# Revealing the combined roles of Aβ and tau in Alzheimer’s disease via a pathophysiological activity decoder

**DOI:** 10.1101/2023.02.21.529377

**Authors:** Lazaro M. Sanchez-Rodriguez, Gleb Bezgin, Felix Carbonell, Joseph Therriault, Jaime Fernandez-Arias, Stijn Servaes, Nesrine Rahmouni, Cecile Tissot, Jenna Stevenson, Thomas K. Karikari, Nicholas J. Ashton, Andréa L. Benedet, Henrik Zetterberg, Kaj Blennow, Gallen Triana-Baltzer, Hartmuth C. Kolb, Pedro Rosa-Neto, Yasser Iturria-Medina

**Affiliations:** Department of Neurology and Neurosurgery, McGill University, Montreal, Canada; McConnell Brain Imaging Centre, Montreal Neurological Institute, Montreal, Canada; Ludmer Centre for Neuroinformatics & Mental Health, Montreal, Canada; McGill University Research Centre for Studies in Aging, Douglas Research Centre, Montreal, Canada; Biospective Inc., Montreal, Canada; Department of Psychiatry and Neurochemistry, Institute of Neuroscience and Physiology, The Sahlgrenska Academy at the University of Gothenburg, Mölndal, Sweden; Department of Psychiatry, School of Medicine, University of Pittsburgh, Pittsburgh, PA, USA; King’s College London, Institute of Psychiatry, Psychology and Neuroscience Maurice Wohl Institute Clinical Neuroscience Institute London UK; NIHR Biomedical Research Centre for Mental Health and Biomedical Research Unit for Dementia at South London and Maudsley NHS Foundation London UK; Centre for Age-Related Medicine, Stavanger University Hospital, Stavanger, Norway; Department of Neurodegenerative Disease, UCL Institute of Neurology, Queen Square, London, UK; UK Dementia Research Institute at UCL, London, UK; Hong Kong Center for Neurodegenerative Diseases, Clear Water Bay, Hong Kong, China; Wisconsin Alzheimer’s Disease Research Center, University of Wisconsin School of Medicine and Public Health, University of Wisconsin-Madison, Madison, WI, USA; Clinical Neurochemistry Laboratory, Sahlgrenska University Hospital, Mölndal; Neuroscience Biomarkers, Janssen Research & Development, La Jolla, California, USA

## Abstract

Neuronal dysfunction and cognitive deterioration in Alzheimer’s disease (AD) are likely caused by multiple pathophysiological factors. However, evidence in humans remains scarce, necessitating improved non-invasive techniques and integrative mechanistic models. Here, we introduce personalized brain activity models incorporating functional MRI, amyloid-β (Aβ) and tau-PET from AD-related participants (N=132). Within the model assumptions, electrophysiological activity is mediated by toxic protein deposition. Our integrative subject-specific approach uncovers key patho-mechanistic interactions, including synergistic Aβ and tau effects on cognitive impairment and neuronal excitability increases with disease progression. The data-derived neuronal excitability values strongly predict clinically relevant AD plasma biomarker concentrations (p-tau217, p-tau231, p-tau181, GFAP). Furthermore, our results reproduce hallmark AD electrophysiological alterations (theta band activity enhancement and alpha reductions) which occur with Aβ-positivity and after limbic tau involvement. Microglial activation influences on neuronal activity are less definitive, potentially due to neuroimaging limitations in mapping neuroprotective vs detrimental phenotypes. Mechanistic brain activity models can further clarify intricate neurodegenerative processes and accelerate preventive/treatment interventions.

## Introduction

Alzheimer’s disease (AD) is defined by synaptic and neuronal degeneration and loss accompanied by amyloid beta (Aβ) plaques and tau neurofibrillary tangles (NFTs) (Iturria-Medina et al., 2018; Jack et al., 2018; Maestú et al., 2021). *In-vivo* animal experiments indicate that both Aβ and tau pathologies synergistically interact to impair neuronal circuits (Busche & Hyman, 2020). For example, the hypersynchronous epileptiform activity observed in over 60% of AD cases (Vossel et al., 2017) may be generated by surrounding Aβ and/or tau deposition yielding neuronal network hyperactivity (Tok et al., 2022; Vossel et al., 2017). Cortical and hippocampal network hyperexcitability precedes memory impairment in AD models (Kazim et al., 2017; Targa Dias Anastacio et al., 2022). In an apparent feedback loop, endogenous neuronal activity, in turn, regulates Aβ aggregation, in both animal models and computational simulations (Bero et al., 2011; de Haan et al., 2017). Multiple other factors involved in AD pathogenesis -remarkably, neuroinflammatory dysregulations-also seemingly influence neuronal firing and act on hypo/hyperexcitation patterns (Iturria-Medina et al., 2016; Kwon & Koh, 2020; Shen et al., 2018). Thus, mounting evidence suggest that neuronal excitability changes are a key mechanistic event appearing early in AD and a tentative therapeutic target to reverse disease symptoms (Busche & Hyman, 2020; Lauterborn et al., 2021; Maestú et al., 2021; Targa Dias Anastacio et al., 2022). However, the exact patterns of Aβ, tau and other disease factors’ neuronal activity alterations in AD’s neurodegenerative progression are unclear as *in-vivo* and non-invasive measuring of neuronal excitability in human subjects remains impractical.

Brain imaging and electrophysiological monitoring constitute a reliable readout for brain network degeneration likely associating with AD’s neuro-functional alterations (Babiloni et al., 2013; Iturria-Medina et al., 2017; Maestú et al., 2021; Sanchez-Rodriguez et al., 2018; Yang et al., 2018). Patients present distinct resting-state blood-oxygen-level-dependent (BOLD) signal content in the low frequency fluctuations range (0.01–0.08 Hz) (Yang et al., 2018, 2020). These differences increase with disease progression, from cognitively unimpaired (CU) controls to mild cognitive impairment (MCI) to AD, correlating with performance on cognitive tests (Yang et al., 2018). Another characteristic functional change is the slowing of the electro-(magneto-) encephalogram (E/MEG), with the signal shifting towards low frequency bands (Babiloni et al., 2013; Sanchez-Rodriguez et al., 2018). Electrophysiological spectral changes associate with brain atrophy and with losing connections to hub regions including the hippocampus, occipital and posterior areas of the default mode network (Maestú et al., 2019). All these damages are known to occur in parallel with cognitive impairment (Maestú et al., 2019). Disease processes also manifest differently given subject-specific genetic and environmental conditions (Iturria-Medina et al., 2018, 2021). Models of multiple pathological markers and physiology represent a promising avenue for revealing the connection between individual AD fingerprints and cognitive deficits (Maestú et al., 2021; Sanchez-Rodriguez et al., 2018; van Nifterick et al., 2022).

In effect, large-scale neuronal dynamical models of brain re-organization have been used to test disease-specific hypotheses by focusing on the corresponding causal mechanisms (Deco et al., 2018; Luppi et al., 2022; Stefanovski et al., 2019). By considering brain topology (the structural connectome (Sanchez-Rodriguez et al., 2018)) and regional profiles of a pathological agent (Deco et al., 2018), it is possible to recreate how a disorder develops, providing supportive or conflicting evidence on the validity of a hypothesis (Luppi et al., 2022). Generative models follow average activity in relatively large groups of excitatory and inhibitory neurons (neural masses), with large-scale interactions generating E/MEG signals and/or functional MRI observations (Iturria-Medina & Evans, 2021). Through neural mass modeling, personalized virtual brains were built to describe Aβ pathology effects on AD-related EEG slowing (Stefanovski et al., 2019). Simulated resting-state functional MRI across the AD spectrum was used to estimate biophysical parameters associated with cognitive deterioration (Zimmermann et al., 2018). In addition, different intervention strategies to counter neuronal hyperactivity in AD have been tested (de Haan et al., 2017; van Nifterick et al., 2022). Notably, comprehensive computational approaches combining pathophysiological patterns and functional network alterations allow the quantification of non-observable biological parameters (Falcon et al., 2016) like neuronal excitability values in a subject-specific basis (Deco et al., 2018; Iturria-Medina et al., 2018, 2021; Luppi et al., 2022; Maestú et al., 2021; Sanchez-Rodriguez et al., 2018), facilitating the design of personalized treatments targeting the root cause(s) of functional alterations in AD.

Here, we considerably extend previous mechanistic brain models of disease progression in four fundamental ways. First, we develop a personalized whole-brain neural mass model integrating multilevel, multifactorial pathophysiological profiles to clarify their causal impact on neuronal activity alterations. Second, using individual *in-vivo* functional MRI together with Aβ- and tau-positron emission tomography (PET), we infer and quantify the combined impact of these relevant AD pathophysiological factors on neuronal excitability. Third, we investigate the associations between the obtained subject-specific pathophysiological neuronal activity affectations and clinically applicable blood-plasma biomarkers (p-tau217, p-tau231, p-tau181, GFAP) as well as cognitive integrity. Fourth, we reproduce hallmark AD electrophysiological alterations and pinpoint their associated critical toxic protein accumulation stages. Overall, our results expand previous understandings of neuropathological impact on AD, namely the emergence of neuronal hyperactivity (Busche & Hyman, 2020; Lauterborn et al., 2021; Maestú et al., 2021; Targa Dias Anastacio et al., 2022), slowing of the E/MEG signals (Babiloni et al., 2013; Sanchez-Rodriguez et al., 2018) and the existence of synergistic multifactorial interactions (Busche & Hyman, 2020; Iturria-Medina et al., 2018). These findings support the premise of using integrative neural mass models to decode multilevel mechanisms in complex neurological disorders.

## Results

### Modeling pathophysiological impacts on whole-brain neuronal activity

A personalized generative framework to study the combined pathophysiological effect of Aβ and tau on neuronal activity was formulated in terms of a whole-brain neuronal mass model (see Figure 1, *Methods*, *Personalized integrative neuronal activity simulator*). This model assumes that, at each brain region, neuronal excitability is potentially mediated by the local pathophysiological burden, specifically by PET-measured accumulation of Aβ plaques, tau tangles and the combined Aβ and tau deposition (their synergistic interaction). In addition, the neuronal populations interact via nervous fibers, potentially propagating pathophysiological effects across each individual brain’s anatomical connectome (Iturria-Medina et al., 2018; Iturria-Medina & Evans, 2015). Altogether, this framework serves to generate continuous pathophysiologically-mediated neuronal activities, which are transformed into BOLD signals by a hemodynamic-metabolic module. The individual model parameters quantifying the brain-wide subject-specific influence of each neuropathological factor (or their synergistic interaction) on neuronal excitability are identified by maximizing the similarity between the generated and observed BOLD data. These estimated parameters serve to reconstruct hidden electrophysiological signals, neuronal excitability spatial profiles, and to study additive relationships with plasma biomarkers and cognitive integrity.

**Figure 1.**
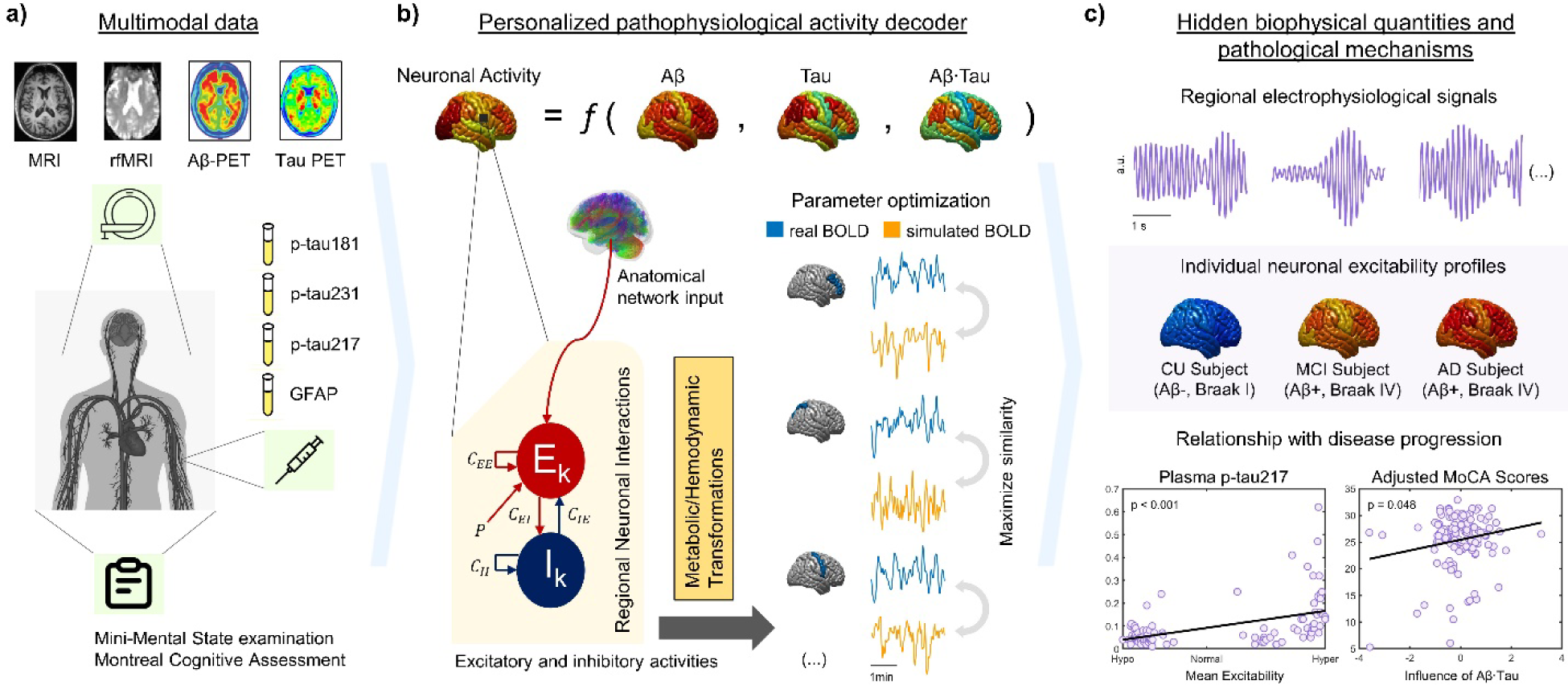
Schematic Personalized Pathophysiological Activity Decoder. (**a**) Individuals underwent a multimodal assessment including structural and resting-state functional MRI, Aβ and tau-PET, clinically relevant plasma biomarkers, and cognitive evaluations. (**b**) In the Alzheimer’s disease model, the subject’s neuronal excitability profile is defined as a function of Aβ, tau and the synergistic interaction of Aβ and tau. Regional excitatory and inhibitory firing rates are influenced by the local pathophysiological profiles and the signals coming from other regions via the anatomical connectome. The regional neuronal signals generate BOLD indicators through metabolic/hemodynamic transformations. By maximizing the similarity between the generated and observed BOLD data, the set of subject-specific influences of the pathophysiological Aβ, tau and Aβ·tau factors on neuronal activity are quantified. (**c**) These estimated pathophysiological influences serve to recover electrophysiological activity producing the real individual BOLD signals, and to study individual excitability profiles and their relationship with independent AD (plasma) markers and cognitive deterioration.

Data from one hundred thirty-two individuals from the Translational Biomarkers in Aging and Dementia cohort (TRIAD, https://triad.tnl-mcgill.com/) were used in the study, including CU (N=81), MCI (N=35) and AD (N=16) participants (Supplementary file 1—table 1). All subjects were cognitively profiled –e.g., MMSE (Folstein et al., 1975), MoCA (Nasreddine et al., 2005)– and underwent structural and resting-state functional MRI and Aβ (^18^F-NAV4694)-, tau (^18^F-MK-6240)- and microglial activation (^11^C-PBR28)-PET. From the fMRI signals, regional fractional amplitudes of low-frequency fluctuations (fALFF) values were obtained, a measure consistently identified as a reliable neuronal activity biomarker of AD’s progression (Iturria-Medina et al., 2018; Yang et al., 2018, 2020). From all the PET images, the corresponding mean Standardized Uptake Value Ratios (SUVRs) (Iturria-Medina et al., 2018; Pascoal et al., 2021; Therriault et al., 2021) were extracted for 66 bilateral regions of interest *(Methods*, *Image processing*). Individuals also had measures of plasma p-tau (Ashton et al., 2021a; Karikari et al., 2020; Therriault, Vermeiren, et al., 2022; Triana-Baltzer et al., 2021) and glial fibrillary acidic protein (Benedet et al., 2021) (*Methods*, *Plasma biomarkers*).

**Table 1.**
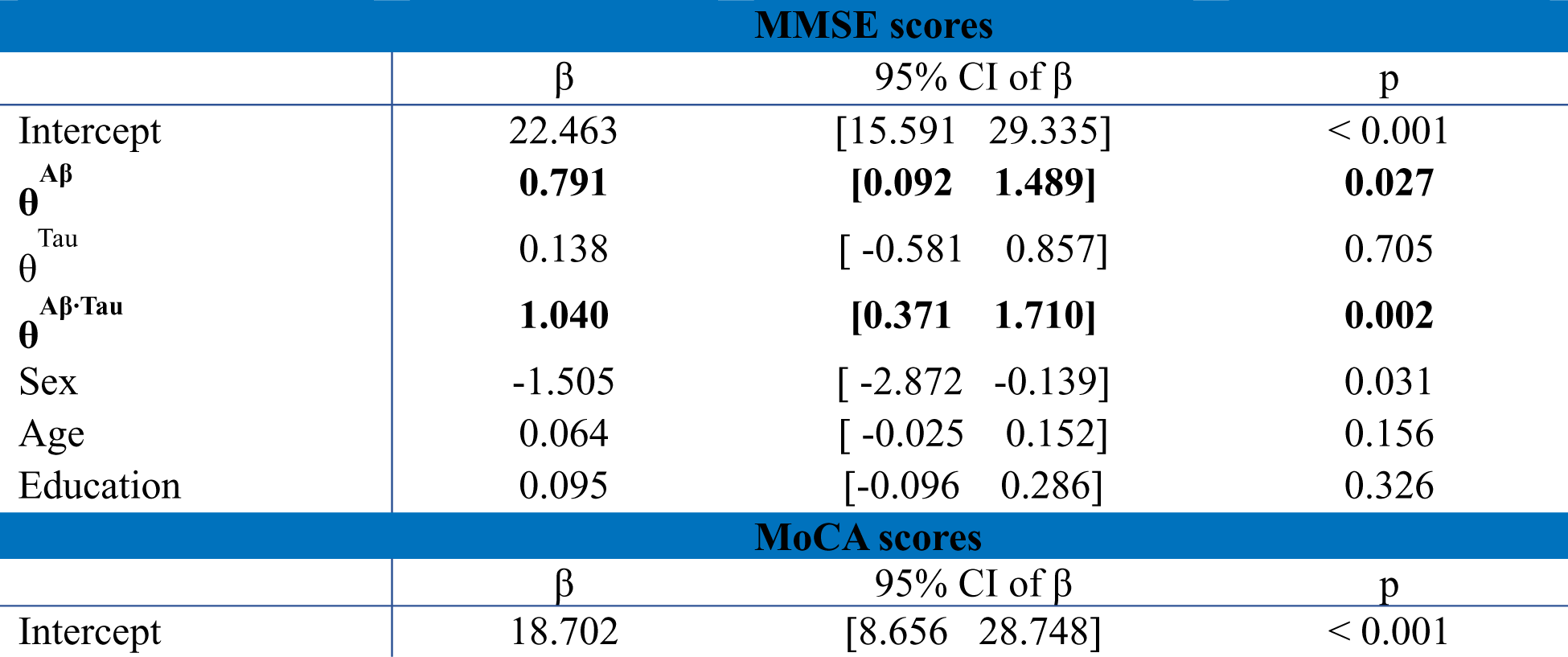
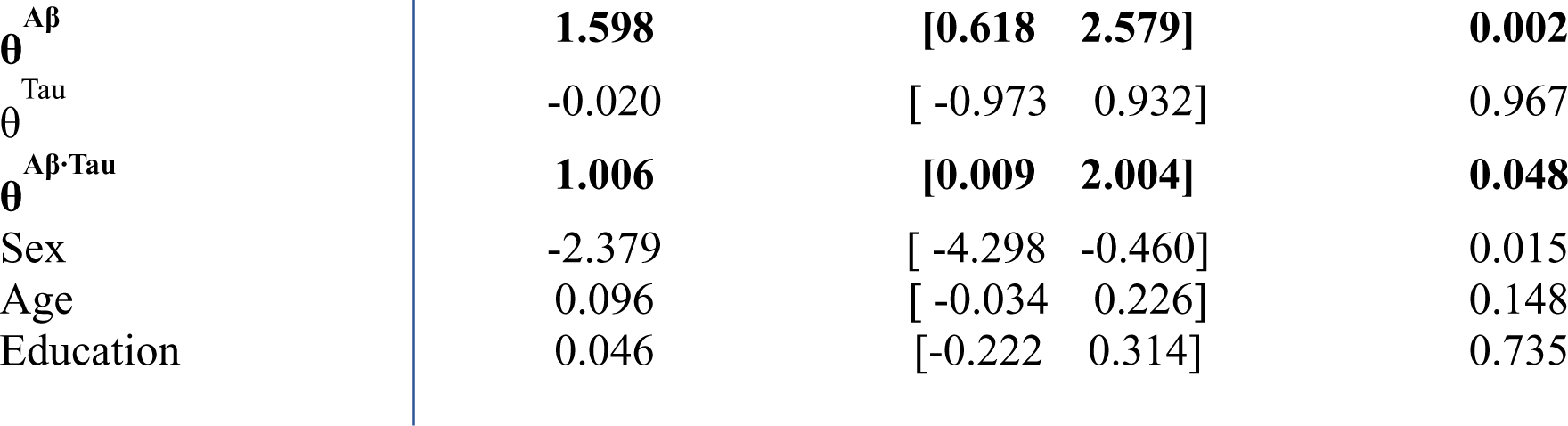
Multiple linear regression analysis investigating the pathological effects on neuronal activity as predictors of MMSE and MoCA scores. The influences of Aβ plaques (θ^Aβ^_E_), tau tangles (θ^Tau^_E_) and the interaction of Aβ and tau (θ^Aβ·Tau^_E_) on neuronal activity, sex, age and education were considered as predictors. Reported values are obtained coefficients (β), the 95% confidence intervals and the p-values for the t-statistic of the two-sided hypothesis tests. Significant pathophysiological terms (5% level) are highlighted. MMSE: R^2^=0.18, p < 0.001; MoCA: R^2^=0.19, p < 0.001. MMSE, Mini-Mental State examination; MoCA, Montreal Cognitive Assessment.

| MMSE scores |  |  |  |  |
| --- | --- | --- | --- | --- |
| | $\beta$ | 95% CI of $\beta$ | | p |
| Intercept | 22.463 | [15.591 | 29.335] | < 0.001 |
| $\theta^{A\beta}$ | **0.791** | **[0.092** | **1.489]** | **0.027** |
| $\theta^{Tau}$ | 0.138 | [-0.581 | 0.857] | 0.705 |
| $\theta^{A\beta \cdot Tau}$ | **1.040** | **[0.371** | **1.710]** | **0.002** |
| Sex | -1.505 | [-2.872 | -0.139] | 0.031 |
| Age | 0.064 | [-0.025 | 0.152] | 0.156 |
| Education | 0.095 | [-0.096 | 0.286] | 0.326 |
| MoCA scores |  |  |  |  |
| | $\beta$ | 95% CI of $\beta$ | | p |
| Intercept | 18.702 | [8.656 | 28.748] | < 0.001 |

|  |  |  |  |
| --- | --- | --- | --- |
| $\theta^{A\beta}$ | **1.598** | **[0.618 2.579]** | **0.002** |
| $\theta^{Tau}$ | -0.020 | [-0.973 0.932] | 0.967 |
| $\theta^{A\beta \cdot Tau}$ | **1.006** | **[0.009 2.004]** | **0.048** |
| Sex | -2.379 | [-4.298 -0.460] | 0.015 |
| Age | 0.096 | [-0.034 0.226] | 0.148 |
| Education | 0.046 | [-0.222 0.314] | 0.735 |

### Reproducing hallmark electrophysiological alterations in AD progression

A desired attribute of biologically-defined generative tools in clinical applications is to reproduce and mechanistically clarify reported pathophysiological observations. We obtained subject-specific relative contributions of the considered pathophysiological factors on neuronal activity (Supplementary file 1—figure 1) and reconstructed proxy quantities for electro-(magneto)encephalographic (E/MEG) sources in each brain region. Subsequently, we aimed to test the proposed pathophysiological activity generator’s ability to recreate reported spectral changes in AD, i.e., increases of theta band power (4–8 Hz) and decreases of power in the lower alpha band (alpha1, 8–10 Hz) (Babiloni et al., 2013; Sanchez-Rodriguez et al., 2018; van Nifterick et al., 2022). Among the quantities contributing to the E/MEG model output, we also closely studied excitatory firings and changes to their magnitude given the influence of the toxic protein depositions (*Methods, Personalized integrative neuronal activity simulator*). The participants were, after individual parameter identification, separated into groups (Supplementary file 1—table 2) according to their clinical diagnosis (CU, MCI, AD) and Aβ-positivity or *in-vivo* Braak staging (Braak & Braak, 1991; Therriault, Pascoal, et al., 2022). We performed statistical tests on the reconstructed quantities of interest to understand the generalized Aβ and tau effects on neuronal activity –see also *Methods, Statistical analyses*, and Supplementary file 1—table 3.

We observed that the standardized ratio of power in the theta band (4–8 Hz) was higher for Aβ+ groups than for Aβ-(Figure 2a). Conversely, the alpha1 (8–10 Hz) power decreased with Aβ-positivity. Finally, the average excitatory firings were generally higher for Aβ+ subjects. Similar results were observed across Braak stages (Figure 2b). Differences between all, theta and alpha1 power and mean excitatory activity, were observed for subjects in Braak 0 (non-significant tau neurofibrillary tangle involvement) and the advanced limbic (Braak III-IV) and isocortical stages (Braak V-VI) and, furthermore, for Braak I-II (transentorhinal) and Braak V-VI subjects.

**Figure 2.**
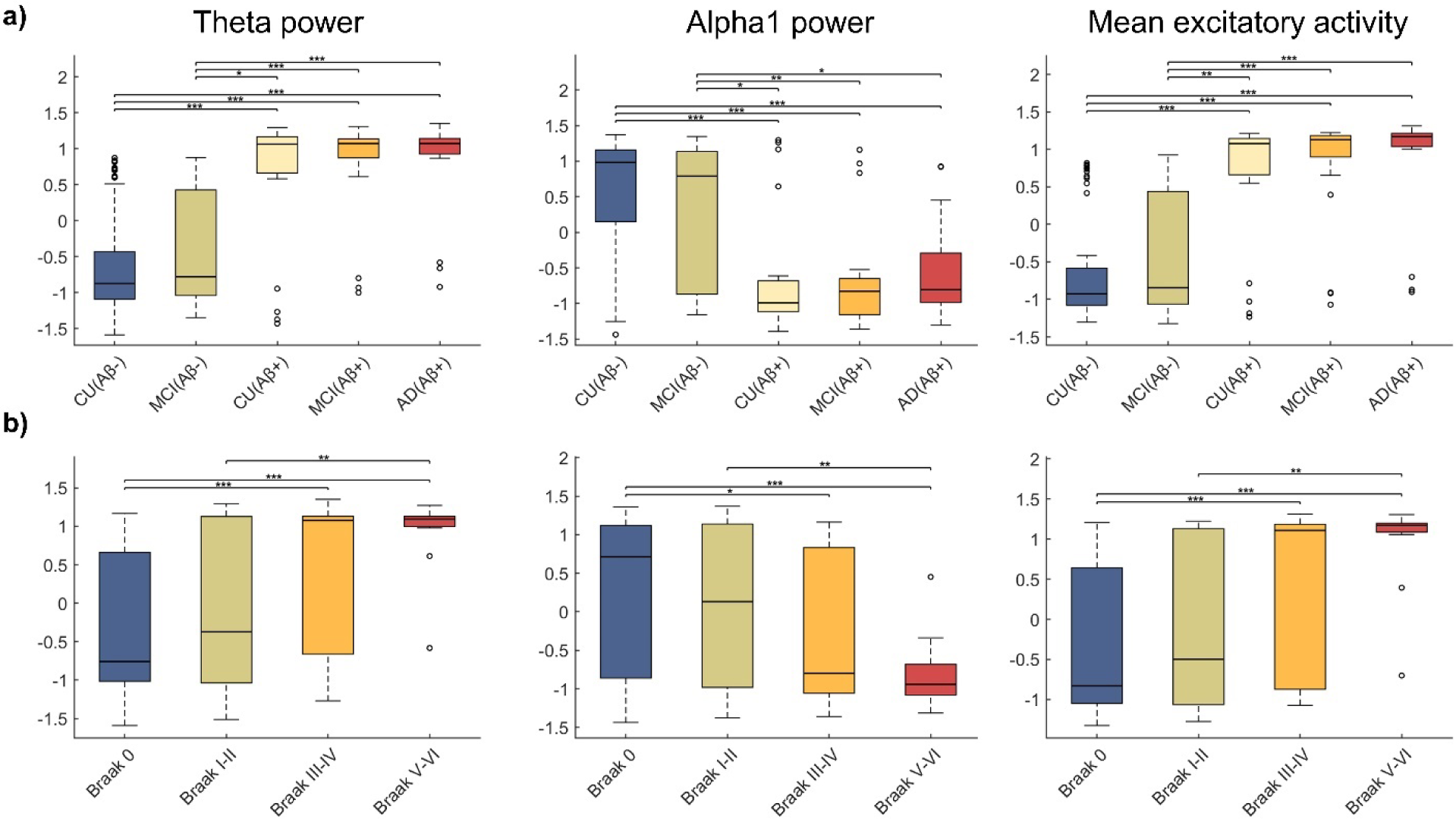
Electrophysiological impacts by *Aβ, tau, Aβ·tau*. From left to right: ratio of power in the theta band (4–8 Hz) of the regional excitatory input currents (the E/MEG is proportional to the excitatory input current), ratio of power in the alpha1 band (8–10 Hz) and mean excitatory firings (over all regions and time points). Each of the quantities was standardized using the mean and s.d. from all subjects, for visualizing general trends. Participants were then grouped according to clinical diagnosis and (**a**) Aβ-positivity and (**b**) Braak stages. In the box-and-whisker plot, the central lines indicate the group medians, with the bottom and top edges of each box denoting the 25th and 75th percentiles, respectively. Whiskers extend to the maximum and minimum values while data points that are deemed outliers for the group are plotted individually with circles. The results of ANCOVA post-hoc t-tests for the above-mentioned groups, with the corresponding electrophysiological quantity as response variable and age and sex as covariates are also shown. * represents significance level p < 0.05, ** means significance level p < 0.01 and *** is p < 0.001.

### Differences in neuronal excitability associate with clinical states and disease progression

Seeking to find mechanisms underlying the observed electrophysiological patterns, we reconstructed the biophysical quantity that changes due to the influence of the pathophysiological factors in our model: neuronal excitability. Figure 3 and Figure 3—figure supplement 1 show excitability values for all brain regions of interest and subjects. The combined action of the pathological factors either increases (“hyper”) or decreases (“hypo”) regional excitability around a certain baseline normal value.

**Figure 3.**
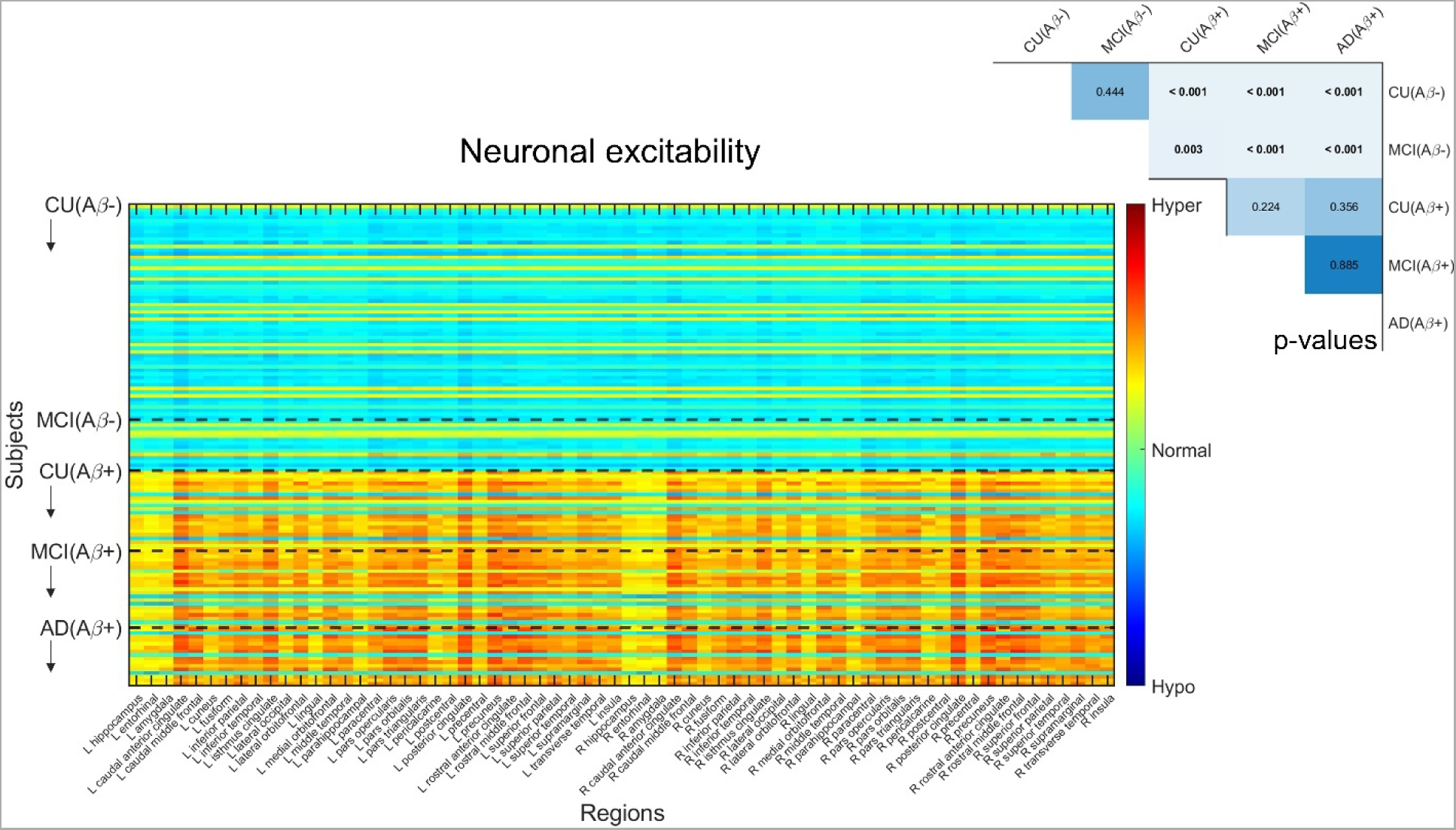
Neuronal excitabilities under the influence of *Aβ, tau and Aβ·tau*. Inferred neuronal excitability values for the brain regions of interest (“y”-axis) and all subjects (“x”-axis). Participants were grouped according to clinical diagnosis and Aβ-positivity in this figure, to understand Aβ’s contribution to the individually estimated biological profiles. Within a group, subjects appear according to their existing ordering in the anonymized database. Warm colors represent hyperexcitability of the region in the subject’s brain and cool colors denote hypoexcitable states. Results of ANCOVA post-hoc t-tests for the above-mentioned groups, with the average intra-brain excitability values as response variable and age and sex as covariates appear in the upper right. P-values in bold fonts represent differences at a 5% significance level or lower.

We found significant differences of neuronal excitability due to Aβ positivity and Braak stages. Firstly, we observed significant discrimination between all Aβ- and Aβ+ groups (Figure 3), i.e.: CU(Aβ-) and CU(Aβ+), MCI(Aβ+), AD(Aβ+) (p < 0.001, sex and age adjusted); MCI(Aβ-) and CU(Aβ+), MCI(Aβ+), AD(Aβ+) (p < 0.05, sex and age adjusted). Additionally, we discovered similar differences between Braak 0 participants and those in all later stages, and for Braak I-II, and Braak V-VI (Figure 3—figure supplement 1). Subjects in advanced disease stages generally presented hyperexcitability profiles, while most of the Aβ- and Braak 0, I-II participants were largely characterized by a slight hypoexcitability.

### Neuronal hyperexcitability relates to high levels of plasma AD biomarkers

In this section, we investigated the relationship between the obtained individual excitability values and blood biomarkers of AD pathophysiology, which constitute accessible alternatives to neuroimaging indicators (Ashton et al., 2021b; Benedet et al., 2021; Therriault, Vermeiren, et al., 2022; Tissot et al., 2021). Figure 4 shows the relationships between the average intra-brain excitabilities and the plasma biomarkers p-tau181, p-tau231 and p-tau217 (phosphorylated tau indicators) and glial fibrillary acidic protein –GFAP, a measure of reactive astrogliosis and neuronal damage (Benedet et al., 2021). Notably, we observed that high levels of the plasma biomarkers significantly relate to the participants’ neuronal hyperactivation.

**Figure 4.**
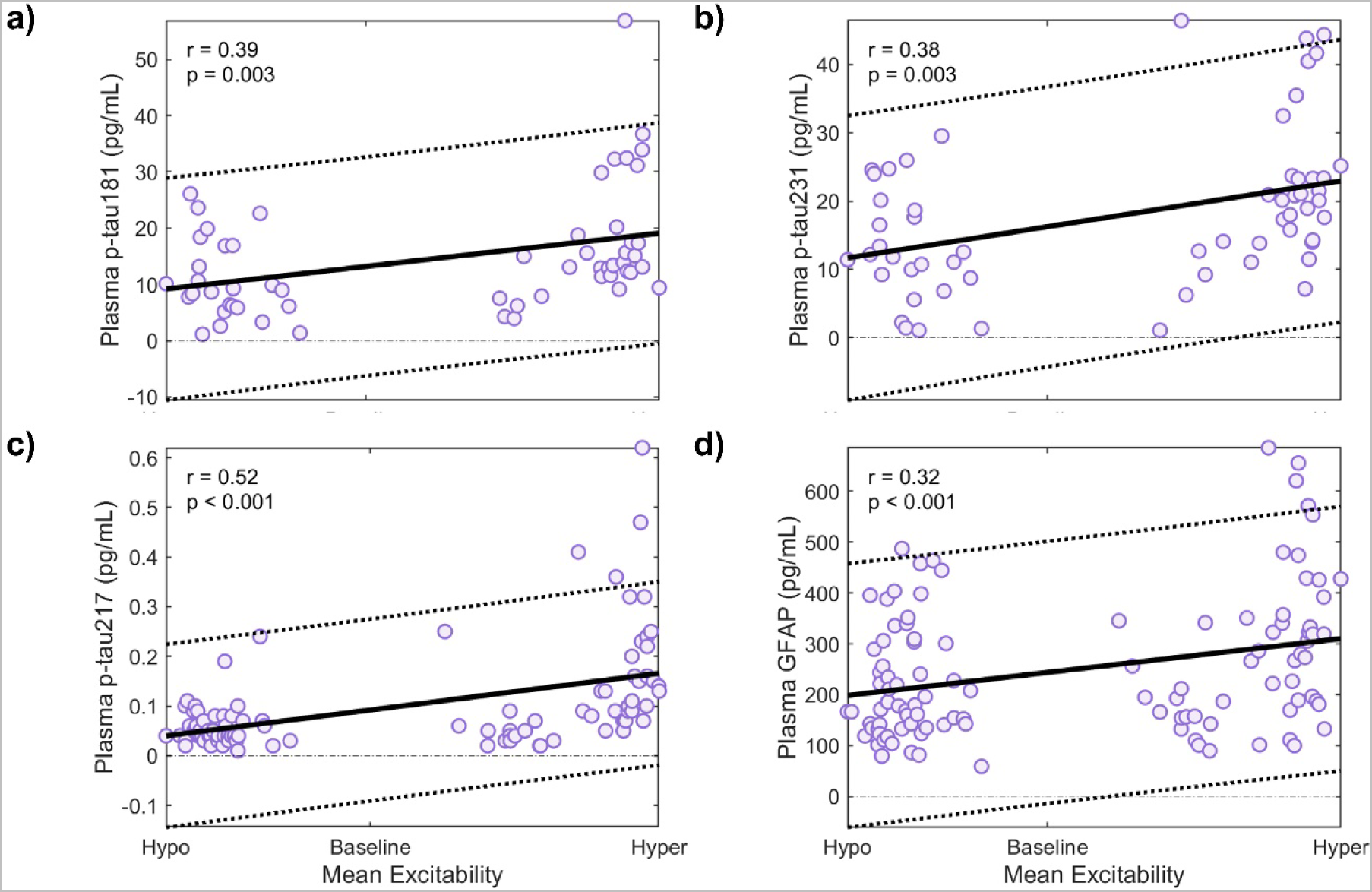
Neuronal excitabilities and AD plasma biomarkers. Spearman’s correlation analyses between the participants’ plasma biomarkers and their estimated average intra-brain excitabilities, for (**a**) p-tau181, (**b**) p-tau231, (**c**) p-tau217 and (**d**) GFAP. The error bands denote 95% confidence intervals (CIs).

### Synergistic Aβ and tau interaction strongly relates to cognitive performance

Finally, we proceeded to estimate the pathophysiological factors’ effects on cognitive impairment through the changes in neuronal activity, measured with the individual weights contributing to neuronal excitability. Table 1 shows the results of regression analyses taking MMSE and MoCA scores as response variables, and the obtained personalized models’ influences of Aβ, tau and the Aβ·tau interaction as predictors, while adjusting for sex, age and education (*Methods, Statistical analyses*). We observed that both the Aβ’s solo influence on neuronal activity and the Aβ·tau synergistic interaction term were significant predictors of MMSE and MoCA evaluations (p < 0.05). The coefficients of these terms in the linear models were positive in all cases. Thus, highly negative influences of the studied pathophysiological factors –yielding hyperexcitability in the model (*Methods, Personalized integrative neuronal activity simulator)*– are linked to low cognitive scores.

## Discussion

We developed an integrative biophysical framework to map pathophysiological influences on neuronal activity, with application to AD. Previous work has investigated how pathological electrophysiological activity emerges in generative models that consider the influence of isolated biological factors, such as Aβ plaques (Stefanovski et al., 2019) or in several possible AD synaptic dysfunction scenarios (de Haan et al., 2017; van Nifterick et al., 2022). Despite the high computational value of these works, realistic biological information could have been estimated from the data under certain constraints, to validate the mechanistic simulations. On the other hand, highly data-driven models (not intended to replicate neuronal activity features) were used to individually characterize multifactorial dynamic interactions propagating through anatomical and vascular networks in the AD spectrum (Adewale et al., 2021; Iturria-Medina et al., 2017, 2018; Khan et al., 2022). Building on the strengths of both mechanistic and data-driven models, our hybrid approach decodes, for the first time, the subject-specific simultaneous and combined pathological neuronal activity contributions of relevant disease factors: Aβ and tau, evaluated through individual functional MRI and PET biomarkers. Altogether, we observed increased neuronal excitability with AD progression, which also predicted increased plasma biomarkers concentrations and cognitive impairment.

Our findings confirm previous observations (Babiloni et al., 2013; Busche & Hyman, 2020; Maestú et al., 2021; Targa Dias Anastacio et al., 2022; Tok et al., 2022; Vossel et al., 2017) and cast new light on microscopical pathological processes that are inaccessible to traditional neuroimaging methods. Through considering the influence of multiple pathophysiological factors, we have retrieved the AD electrophysiological hallmark: enhancement of theta band activity together with alpha decreases, as disease progresses (Babiloni et al., 2013; Sanchez-Rodriguez et al., 2018) from BOLD signals. Our results also indicate that CU(Aβ+) and/or Braak III-IV are the stages from which these electrophysiological biomarkers become abnormal. These groups contain subjects who are not cognitively impaired but present significant Aβ deposition (Therriault et al., 2021) and/or have widespread temporal and parietal tau aggregation detectable by tau PET (Braak & Braak, 1991). A recent study, also on subjects from the TRIAD cohort, found reduced, clinically significant delayed recall and recognition memory tests performance at Braak III and IV stages as well (Fernández Arias et al., 2023). Additionally, multicenter research has shown that CU(Aβ+) subjects, independently of tau status, present substantially increased risk of short-term (3-5 years) conversion to mild cognitive impairment, compared to CU(Aβ-) (Ossenkoppele et al., 2022). Our observations reaffirm this evidence. Aβ+ and post-Braak II individuals may be the most likely candidates to benefit from early disease interventions modifying the cognitive decline that associates with patho-electrophysiological activity (Babiloni et al., 2013; Maestú et al., 2019; Sanchez-Rodriguez et al., 2018).

No previous direct *in-vivo* evidence for AD-associated neuronal hyperactivity existed thus far in humans, although proxy measurements (Celone et al., 2006), post-mortem studies (Lauterborn et al., 2021) and animal models (Busche & Hyman, 2020) have suggested a similar mechanism. In this study, by assuming a toxic protein influence model (*Aβ, tau, Aβ·tau*) we inferred neuronal excitability values from the individual PET-functional MRI datasets. Given the experimental conditions (equal nominal parameters, anatomical connectivities and model assumptions), the estimation of the pathophysiological influences is totally unsupervised with each subject’s model being blind to the others’ imaging data and clinical assessments. As such, the broad association of subjects in advanced disease stages with hyperactivity –and conversely– is only driven by the individual PET datasets and the maximization of the similarity between the real resting-state functional MRI and the simulated BOLD signals. The progression towards hyperexcitation with disease worsening was equally evident for a simplified model with separate contributions by Aβ and tau only (Supplementary file 1—figures 2-3). Increased excitability was also associated with high levels of plasma biomarkers (blood phosphorylated tau and GFAP) which are sensitive to incipient AD pathology (Ashton et al., 2021b, 2022; Benedet et al., 2021; Milà-Alomà et al., 2022; Tissot et al., 2021) and disease progression, especially p-tau217 (Ashton et al., 2021b; Therriault, Vermeiren, et al., 2022), further demonstrating the strong biomarker capabilities of our estimated excitability parameters. Additionally, we observed that the more hyperactive the existing excitatory neuronal populations of a subject were (given by negative influence values of the significant factors in our model), the greater the participant’s cognitive dysfunction, thus supporting a direct link among neuronal excitability, pathophysiological burden, and cognitive integrity.

Beyond AD-related protein deposition, our method can also investigate the influence of other critical factors. It has been hypothesized, and to some extent observed (Kwon & Koh, 2020; Shen et al., 2018; Targa Dias Anastacio et al., 2022), that microglial activation, a probable marker for neuroinflammation (Nutma et al., n.d.-a; Pascoal et al., 2021), affects excitability and neuronal activity in AD. Consequently, we performed a set of complimentary experiments where we recreated the obtained results in a model that also considers deviations to neuronal excitability due to microglial activation –measured with 18kDa Translocator Protein PET. However, we did not find separation between clinical groups in terms of the estimated neuronal excitabilities when the microglial activation factor was considered (Supplementary file 1—figures 4-5). Moreover, the synergistic interaction of Aβ and tau was the factor that better predicted cognitive impairment, with no significant effect by the microglial activation term (Supplementary file 1—table 4). We attribute this effect to technical limitations associated with the acquisition of microglial activation. Unlike the Aβ and tau PET SUVRs data, which showed extended statistically significant differences across all brain regions for CU and AD participants (two-sampled t-test, p < 0.05), microglial activation images exhibited differences in only 24 regions (i.e., approx. 36%; Supplementary file 1—table 5). Microglial activation is thought to have a neuroprotective character (M2-phenotype) at early disease stages (Kwon & Koh, 2020; Shen et al., 2018). On the other hand, excessive activation of microglia seemingly becomes detrimental in clinical AD (M1-phenotype) by releasing pro-inflammatory cytokines that may exacerbate AD progression (Kwon & Koh, 2020; Pascoal et al., 2021; Shen et al., 2018). Nevertheless, modern neuroinflammation PET tracers are not specific to these two different phenotypes as no consistent targets have been discovered (Shen et al., 2018). Thus, our extended results albeit being relatively uninformative in terms of AD-affectations to neuronal excitability, capture intrinsic microglial activation PET mapping insufficiencies (Nutma et al., n.d.-b).

Our methodology also has limitations. We provide partial validation for the results of the *in-silico*, data-informed approach in the context of the literature only, by reconstructing functional network quantities with well-documented AD affectations for *a posteriori* analysis and by studying relationships with reliable markers of disease progression independently collected in our cohort subjects (plasma concentrations, cognitive scores). This strategy for analyzing the inferred values stems from the lack of ground-truth *in-vivo* excitability measurements. Although we used state-of-the-art fMRI experiments in this study (TR = 681ms, spatial resolution = 2.5×2.5×2.5 mm^3^), more detailed spatiotemporal dynamics could be captured with novel ultra high-resolution functional imaging techniques (Tan Toi et al., n.d.). On the other hand, by using average anatomical connectivity, we have singled-out the mechanisms by which toxic protein deposition and neuroinflammation are associated with pathological neuronal activity. Personalized therapeutic interventions (Iturria-Medina et al., 2018) would require precise individual profiles for increased efficiency. In such applications, including the connectomes’ individual variability may be beneficial. Regarding the neuro-physical model for the influence of pathophysiological factors, two aspects should be considered in future work. Firstly, extending the intra-regional neuronal interactions with additional excitatory and inhibitory populations, pursuing a finer descriptive scale, will also enable us to account for additional significant disease factors such as neuronal atrophy (Iturria-Medina et al., 2016). Secondly, the effects on inhibitory firings should be explored separately as well. Pyramidal (excitatory) neurons greatly outnumber any other neuronal population, making them the most likely proteinopathies target (Maestú et al., 2021). However, inhibitory populations are key in maintaining healthy firing balances (Maestú et al., 2021) and interacting with glial cells (Mederos & Perea, 2019). Finally, the focus of this study was limited to capturing abnormalities in AD by Aβ and tau’s combined action. The model inputs will require modifications to measure neuronal excitability contributions in other neurodegenerative conditions given their characteristic neuropathological factors. For example, dopamine transporter (DaT) ^123^I–FP-CIT scans can be used to quantify dopaminergic deficiency consistent with Parkinsonism and associated disorders (Nichols et al., 2018). Ongoing efforts pursue developing alpha-synuclein protein PET radiotracers that do not also bind to Aβ (Roshanbin et al., 2022). Replacing the AD-pathophysiology with such quantified maps in our framework may well help advance the characterization of neuronal excitability dysfunction in the Parkinsonian circuit (Picconi et al., 2012).

Our approach has major implications to disease hypothesis testing. Generative models (Luppi et al., 2022) in works by Iturria-Medina et al. (Iturria-Medina et al., 2017, 2018), Deco et al. (Deco et al., 2018, 2021), Sotero et al. (Sanchez-Rodriguez et al., 2018; Sotero & Trujillo-Barreto, 2008), de Haan et al. (de Haan et al., 2017; van Nifterick et al., 2022) among others, focus on better comprehending neurological conditions. The models considered in the present study reflect plausible biophysical mechanisms determining neuronal activity abnormalities in the AD spectrum (Busche & Hyman, 2020; Kwon & Koh, 2020; Maestú et al., 2021; Targa Dias Anastacio et al., 2022). Critical mechanistic information on the underlying activity-generating processes is obtained, as well as about their relationship with clinical and cognitive profiles, as all these disease-informative variables are tracked in our comprehensive methodology. Importantly, we observed that the synergistic interaction of Aβ and tau, and Aβ separate contributions are the most significant factors influencing aberrant neuronal activity, AD progression and symptomatology. The relative preponderance of Aβ’s effect with respect to tau’s was somewhat expected as Aβ plaques generalize to many cortical areas early in the disease, while NFT spreading increases rapidly in temporal and parietal regions only (Insel et al., 2020). These pathological progression patterns, measured by PET uptake, inform our individual dynamical models. A critical methodological contribution is the capacity to resolve complex biological processes hidden to current non-invasive imaging and electrophysiological monitoring techniques, e.g., the neural masses’ firing excitabilities. For future work, we aim to further clarify the specific molecular features responsible for the differences in excitability values across clinical stages. By doing so, we expect to gain additional insights into AD pathophysiology that could boost diagnostic accuracy and preclinical applications. This *pathophysiological activity decoder* is equally applicable to other intricate multifactorial neurological disorders by considering their relevant disease factors. Computational disease modeling may further unveil the complex mechanisms of neurodegeneration and aid providing efficient treatment at a personalized level.

## Methods

### Participants

We selected individuals from the Translational Biomarkers in Aging and Dementia (TRIAD) cohort (https://triad.tnl-mcgill.com/). The study was approved by the Douglas Mental Institute Research Board and all participants gave written consent. All subjects underwent MRI, resting-state fMRI, Aβ (^18^F-NAV4694)-, tau (^18^F-MK-6240)- and neuroinflammation (^11^C-PBR28)-PET scans, together with a complete cognitive evaluation, including the Mini-Mental State Examination (MMSE) and the Montreal Cognitive Assessment (MoCA). We chose baseline assessments in all cases. Only participants with “cognitively unimpaired” (N=81), “mild cognitive impairment” (N=35), or “probable Alzheimer’s disease” (N=16) clinical and pathophysiological diagnoses were considered (Tissot et al., n.d.).

### Image processing

#### MRI

Brain structural T1-weighted 3D images were acquired for all subjects on a 3 T Siemens Magnetom scanner using a standard head coil. T1 space sequence was performed in sagittal plane in 1 mm isotropic resolution; TE 2.96 ms, TR 2300 ms, slice thickness 1 mm, flip angle 9 deg, FOV read 256 mm, 192 slices per slab. The images were processed following a standard pipeline (Iturria-Medina et al., 2018), namely: non-uniformity correction using the N3 algorithm, segmentation into grey matter, white matter and cerebrospinal fluid (CSF) probabilistic maps (SPM12, www.fil.ion.ucl.ac.uk/spm) and standardization of grey matter segmentations to the MNI space (Evans et al., 1994) using the DARTEL tool (Ashburner, 2007). Each map was modulated to preserve the total amount of signal/tissue. We selected 66 (bilateral) cortical regions in the Desikian-Killiany-Touriner (DKT) (Klein & Tourville, 2012) atlas (Supplementary file 1—table 5). Subcortical regions, e.g., in the basal ganglia, were not considered given their tendency to present PET off-target binding (Vogel et al., 2020; Young et al., 2020).

#### fMRI

The resting-state fMRI acquisition parameters were: Siemens Magnetom Prisma, echo planar imaging, 860 time points, TR = 681 ms, TE = 32.0 ms, flip angle = 50 deg, number of slices = 54, slice thickness = 2.5 mm, spatial resolution = 2.5×2.5×2.5 mm^3^, EPI factor = 88. We applied a minimal preprocessing pipeline (Iturria-Medina et al., 2018) including motion correction and spatial normalization to the MNI space (Evans et al., 1994) using the registration parameters obtained for the structural T1 image, and removal of the linear trend. We calculated the fractional amplitude of low-frequency fluctuations (fALFF) (Yang et al., 2018), a regional proxy indicator for neuronal activity that has shown high sensibility to disease progression. Briefly, we transformed the signals for each voxel to the frequency domain and computed the ratio of the power in the low-frequency range (0.01–0.08 Hz) to that of the entire BOLD frequency range (0– 0.25 Hz) with code from the RESTplus toolbox (Jia et al., 2019). The fALFF values were ultimately averaged over the voxels according to their belonging to brain regions.

#### Diffusion Weighted MRI (DW-MRI)

High angular resolution diffusion imaging (HARDI) data was acquired for N = 128 in the Alzheimer’s Disease Neuroimaging Initiative (ADNI) (adni.loni.usc.edu). The authors obtained approval from the ADNI Data Sharing and Publications Committee for data use and publication, see documents http://adni.loni.usc.edu/wp-content/uploads/how_to_apply/ADNI_Data_Use_Agreement.pdf and http://adni.loni.usc.edu/wp-content/uploads/how_to_apply/ADNI_Data_Use_Agreement.pdf and http://adni.loni.usc.edu/wp-content/uploads/how_to_apply/ADNI_Manuscript_Citations.pdf, respectively. For each diffusion scan, 46 separate images were acquired, with 5 b_0_ images (no diffusion sensitization) and 41 diffusion-weighted images (b = 1000 s/mm^2^). ADNI aligned all raw volumes to the average b_0_ image, corrected head motion and eddy current distortions. Region-to-region anatomical connection density matrices were obtained using a fully automated fiber tractography algorithm (Iturria-Medina et al., 2007) and intravoxel fiber distribution reconstruction (Tournier et al., 2008). For any subject and pair of regions *k* and *l*, the ∁_*lk*_ measure (0 ≤ ∁_*lk*_ ≤ 1, ∁_*lk*_= ∁_*lk*_) reflects the fraction of the region’s surface involved in the axonal connection with respect to the total surface of both regions. More details can be found in a previous publication where ADNI’s DW-MRI was utilized (Iturria-Medina et al., 2018). We averaged the ADNI subject-specific connectivity matrices (Iturria-Medina et al., 2018; Sanchez-Rodriguez et al., 2021) to utilize a single, representative anatomical network across our calculations on the TRIAD dataset.

#### PET

Study participants had Aβ (^18^F-NAV4694), tau (^18^F-MK-6240) and translocator protein microglial activation (^11^C-PBR28) PET imaging in a Siemens high-resolution research tomograph. A bolus injection of ^18^F-NAV4694 was administered to each participant and brain PET imaging scans were acquired approximately 40-70 min post-injection. The images were reconstructed using an ordered subset expectation maximization (OSEM) algorithm on a 4D volume with three frames (3 × 600 s) (Therriault et al., 2021). ^18^F-MK-6240 PET scans of 20 min (4 × 300 s) were acquired at 90-110 min after the intravenous bolus injection of the radiotracer (Pascoal et al., 2020). ^11^C-PBR28 images were acquired at 60–90 min after tracer injection and reconstructed using the OSEM algorithm on a 4D volume with 6 frames (6 × 300 s) (Pascoal et al., 2021). Images were preprocessed according to four main steps (Jagust et al., 2010): 1) dynamic co-registration (separate frames were co-registered to one another lessening the effects of patient motion), 2) across time averaging, 3) re-sampling and reorientation from native space to a standard voxel image grid space (“AC-PC” space), and 4) spatial smoothing to produce images of a uniform isotropic resolution of 8 mm FWHM. Using the registration parameters obtained for the participants’ structural T1 images, all PET images were spatially normalized to the MNI space. ^18^F-MK-6240 images were meninges-striped in native space before performing any transformations to minimize the influence of meningeal spillover. SUVR values for the DKT grey matter regions were calculated using the cerebellar gray matter as the reference region.

The DKT atlas was separately used to define the ROIs for tau-PET Braak stage-segmentation (Braak & Braak, 1991; Therriault, Pascoal, et al., 2022) which consisted of: Braak I (pathology confined to the transentorhinal region of the brain), Braak II (entorhinal and hippocampus), Braak III (amygdala, parahippocampal gyrus, fusiform gyrus and lingual gyrus), Braak IV (insula, inferior temporal, lateral temporal, posterior cingulate and inferior parietal), Braak V (orbitofrontal, superior temporal, inferior frontal, cuneus, anterior cingulate, supramarginal gyrus, lateral occipital, precuneus, superior parietal, superior frontal and rostromedial frontal) and Braak VI (paracentral, postcentral, precentral and pericalcarine) (Braak et al., 1995). All image processing was performed in MATLAB 2021b (The MathWorks Inc., Natick, MA, USA) with the aid of the specific tools and algorithms specified above.

### Plasma biomarkers

Blood biomarkers were quantified with Single molecule array (Simoa) assays (Quanterix, Billerica, MA). These measurements included tau phosphorylated at threonine 181 (p-tau181) (Tissot et al., 2021), tau phosphorylated at threonine 231 (p-tau231) (Ashton et al., 2021b), tau phosphorylated at threonine 217 (p-tau217) (Therriault, Vermeiren, et al., 2022; Triana-Baltzer et al., 2021) and glial fibrillary acidic protein (GFAP) (Benedet et al., 2021) and have been previously reported.

### Personalized integrative neuronal activity simulator

#### Electrophysiological model

The individual electrophysiological brain activity is realized through coupled Wilson-Cowan (WC) modules (Abeysuriya et al., 2018; Daffertshofer & van Wijk, 2011; Gjorgjieva et al., 2016; Meijer et al., 2015; Wilson & Cowan, 1972). In this simple formulation, the variables of interest are the firing rates of the excitatory and inhibitory neural masses, *E*(*t*) and *I*(*t*), respectively. Neural masses are average neuronal populations describing the dynamic behavior of similar neurons within a given spatial domain, i.e., brain regions (Jansen & Rit, 1995; Sanchez-Rodriguez et al., 2018; Wilson & Cowan, 1972). In WC, the excitatory and inhibitory populations are locally coupled. Moreover, the excitatory population is further stimulated by other local inputs (*P*) and cortico-cortical connections, ∁ (*Image processing*, *Diffusion Weighted MRI*).

In effect, each *lk* region influences the dynamics of the *k* region by the quantity 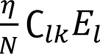, where η is a global scaling coupling strength and *N* is the total number of regions (*N* = 66). We performed a dynamical system analysis (Daffertshofer & van Wijk, 2011; Deco et al., 2009; Gjorgjieva et al., 2016; Wilson & Cowan, 1972) and obtained common *P* and η values simulating plausible electrophysiological oscillations and BOLD signals (Supplementary file 1—figure 6a) for all participants. The parameters used in this study are reported in Supplementary file 1—table 6.

All the inputs (both local and external) received by a neuron are integrated in time when their sum surpasses a certain threshold, (θ_*E*_ or θ_*I*_) (Daffertshofer & van Wijk, 2011). In the neural mass framework, this integration is achieved by means of a sigmoidal activation function (Wilson & Cowan, 1972), 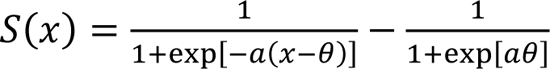. Compared to the “baseline” firings obtained by using the canonical values reported in the literature, regional excitability can be higher (hyperexcitability) or lower (hypoexcitability) depending on whether the firing rate function is shifted to lower or higher input current values, respectively (Supplementary file 1—figure 6b). In our approach, the regional activity profiles are determined by the excitatory firing, a simplification based on the much larger excitatory prevalence in the cortex (Lauterborn et al., 2021; Maestú et al., 2021). We suppose that the regional excitatory firing thresholds are mediated by the following disease factors: Aβ plaques (with a subject-specific contribution weight given by θ^β^), tau tangles (θ^Tau^_E_) and the interaction of amyloid and tau (θ^Aβ.Tau^_E_). These effects are simplistically written as linear fluctuations from the normal baseline value due to the participant’s regional accumulation of each factor (Supplementary file 1—figure 6c), with the SUVRs normalized to the [0,1] interval (Supplementary file 1—figure 7), to preserve the dynamical properties of the desired solution:

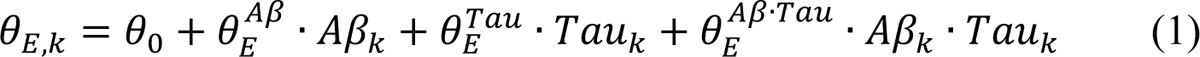

A negative contribution by a factor (θ^Aβ^_E_, θ^Tau^_E_ or θ^Aβ·Tau^_E_) means that the pathological accumulation of such a biomarker tends to decrease the firing threshold thus yielding hyperexcitability. Given the inverse relationship existing between firing thresholds and effective firing rates, we define regional excitability as 1/θ_E,k_.

The evolution of the average firing rates *E*(*tt*) and *I*(*t*) is then given by the following set of differential equations (Daffertshofer & van Wijk, 2011; Gjorgjieva et al., 2016; Wilson & Cowan, 1972):

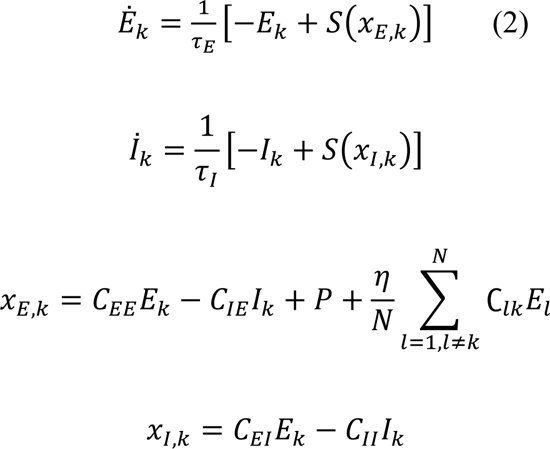

Here, the terms *x*_*E, k*_ and *x*_*I*,*lk*_ are known as the input currents. A synthetic EEG signal is proportional to the regional excitatory input current (Meijer et al., 2015). Since the BOLD signal is related to the afferent neuronal input (Logothetis et al., 2001), we utilized the total action potential arriving to the neuronal populations from other local and external populations as the proxy electrophysiological quantities of interest feeding the metabolic/hemodynamic model by Sotero et al. (Sotero & Trujillo-Barreto, 2008; Valdes-Sosa et al., 2009).

#### Metabolic/hemodynamic model

This biophysical model reflects the role that excitatory and inhibitory activities play in generating the BOLD signal (Sotero et al., 2009; Sotero & Trujillo-Barreto, 2007, 2008; Valdes-Sosa et al., 2009). All variables are normalized to baseline values. Changes in glucose consumption (*g*_*E, lk*_ and *g*_*I,lk*_) are linked to the excitatory and inhibitory (ξ_*E, lk*_ and ξ_*I,lk*_) neuronal inputs in region *k*:

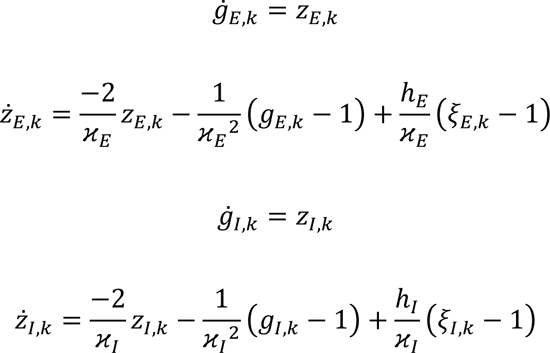

The metabolic rates of oxygen for excitatory (*m*_*E, lk*_) and inhibitory (*m*_*I,lk*_) activities, and the total oxygen consumption, *mm*_*l*_, are obtained from the glucose variables.

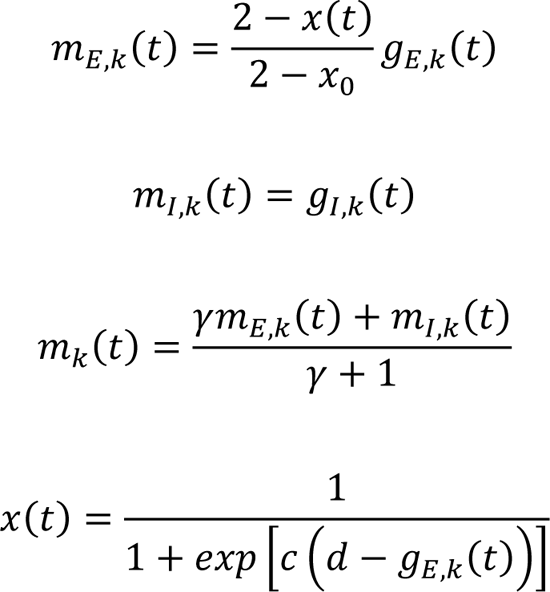

Next, CBF dynamics (*f*_*k*_) is modeled as follows (Friston et al., 2000), assuming that CBF is coupled to excitatory activity:

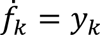

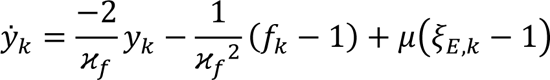

The outputs of the metabolic and vascular models are converted to normalized cerebral blood volume (*b*_*k*_) and deoxy-hemoglobin (*q*_*l*_) content through the Balloon model (Buxton et al., 1998):

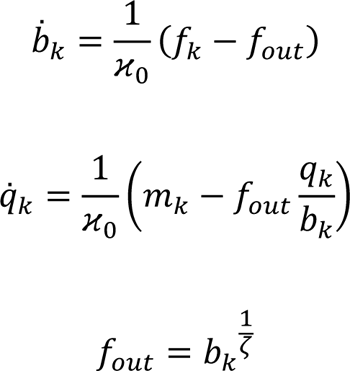

The BOLD signal is obtained by using a linear observation equation as in:

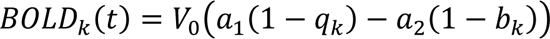

where *a*_1_ = 4.3γ_0_*E*_0_ · *aTE* + εr_0_*E*_0_ ·*aE* and *a*_2_ = εr_0_*E*_0_ · *aE* + ε − 1 are parameters that depend on the experimental conditions (field strength, *aE*) (Archila-Meléndez et al., 2020; Deco et al., 2018; Obata et al., 2004; Simon & Buxton, 2015). The interpretation and specific parameter values of the metabolic/hemodynamic transformations resulting in the simulated BOLD signal can be found in Supplementary file 1—table 7. The sets of equations above were solved, for each individual dataset, with an explicit Runge-Kutta (4,5) method, ode45, as implemented in MATLAB 2021b (The MathWorks Inc., Natick, MA, USA) and a timestep of 0.001s.

#### Parameter estimation

For each participant, we compute the regional fALFF values of the simulated BOLD signals and maximize their similarity (correlation) with the subject’s real BOLD indicators (Supplementary file 1—figure 6d). The estimation of the optimal pathological influences set (θ^Aβ^_E_, θ^*Tau*^_E_, θ^Aβ·*Tau*^_E_) was performed via surrogate optimization (MATLAB 2021b’s surrogateopt). This parameter optimization method performs few objective function evaluations hence it is well-suited for expensive functions as it is the case of our high-dimensional BOLD-simulating dynamical system. We constrained the pathological influences on a small interval around the nominal parameter value (Supplementary file 1—table 6) (Abeysuriya et al., 2018; Daffertshofer & van Wijk, 2011; Gjorgjieva et al., 2016; Meijer et al., 2015; Wilson & Cowan, 1972) to compare results across subjects and disease states. Then, we performed optimization iterations until no new feasible points were found in the allowed interval through 20 sequences of random sample points, guaranteeing that the global minimum was attained (Supplementary file 1—pseudocode 1).

#### Interpreting the pathophysiological effects on neuronal activity

The obtained pathological influences (θ^Aβ^_E_, θ^*Tau*^_E_, θ^Aβ·*Tau*^_E_) describe subject-specific interactions determining brain activity. We use these weights to reconstruct otherwise hidden electrophysiological quantities of interest. Individual neuronal excitability patterns (van Nifterick et al., 2022) are mapped through equation (1) and can be related to separate measurements like plasma biomarkers for AD (Ashton et al., 2021b; Benedet et al., 2021; Tissot et al., 2021). Grand average excitatory activities are found by averaging the firing rates *E*_*lk*_(*t*) over the regions and time points (van Nifterick et al., 2022), for every subject. Likewise, the input currents of equation (2) are used as proxy measures for cortical sources of resting-state EEG (Meijer et al., 2015). We perform a Fast Fourier Transformation power analysis of the neural masses’ signals and determine the relative power of the traditional rhythms, in particular: theta (4–8 Hz) and alpha 1 (8–10 Hz) frequency band oscillations (van Nifterick et al., 2022). Additionally, we investigate the relationship of the obtained pathophysiological influences with cognition (Folstein et al., 1975; Nasreddine et al., 2005).

### Statistical analyses

Clinical diagnosis and PET-imaging Aβ status (determined visually by consensus of two neurologists blinded to the diagnosis) were used to divide the cohort for analyses of the results. Separately, we employed another division based on the conventional unambiguous Braak grouping (Braak & Braak, 1991) of I-II (transentorhinal stages), III–IV (limbic) and V–VI (isocortical), to assess trends in terms of intracellular tau neurofibrillary changes. Group-differences in the electrophysiological quantities of interest (average intra-brain theta and alpha1 power, excitatory firing activity and excitability) were evaluated with ANCOVA post-hoc t-tests, i.e., we looked at the effects of the clinical groups and Aβ positivity/Braak stages on the corresponding quantity, accounting for age and sex. The average theta and alpha1 power and excitatory firing activity were box-cox and z-score transformed across subjects. The associations between excitability and plasma biomarkers were tested using Spearman’s Rho correlation (large-sample approximation). In addition, to assess the relationship between the pathophysiological factors and cognitive integrity we fitted multiple linear regression models using the following specifications: 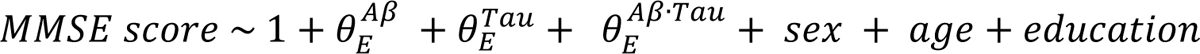 and 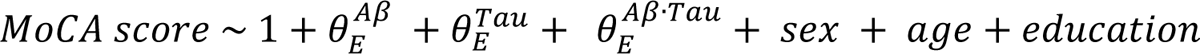 were standardized using the mean and s.d. from all subjects.

## Data availability

The data that support the findings of this study are available by submitting a data share request via https://triad.tnl-mcgill.com/contact-us/. Data are not publicly available due to them containing information that could compromise research participant privacy/consent. All the data collected under the TRIAD cohort is governed by the policies set by the Research Ethics Board Office of the McGill University, Montreal and the Douglas Research Center, Verdun.

## Code availability

The code utilized in this article for the neuronal activity simulations and quantification of the pathological effects conforms to the Open Source Definition and will be freely available with publication at the *Neuroinformatics for Personalized Medicine* lab’s website (NeuroPM, https://www.neuropm-lab.com/publication-codes.html).

## Acknowledgments

LSR was partially supported by funding from the Fonds de recherche du Québec – Santé and the Healthy Brains for Healthy Lives (HBHL) initiative. This project was undertaken thanks in part to the following funding awarded to YIM: the Canada Research Chair tier-2, the CIHR Project Grant 2020, and the Weston Family Foundation’s AD Rapid Response 2018 and Transformational Research in AD 2020. In addition, we used the computational infrastructure of the McConnell Brain Imaging Center at the Montreal Neurological Institute, supported in part by the *Brain Canada Foundation*, through the *Canada Brain Research Fund*, with the financial support of *Health Canada* and sponsors. PRN, GB, JT and JF are supported by the Canadian Institutes of Health Research (CIHR) [MOP-11-51-31; RFN 152985, 159815, 162303], Canadian Consortium of Neurodegeneration and Aging (CCNA; MOP-11-51-31 - team 1), Weston Brain Institute, the Alzheimer’s Association [NIRG-12-92090, NIRP-12-259245], Brain Canada Foundation (CFI Project 34874; 33397), the Fonds de Recherche du Québec – Santé (FRQS; Chercheur Boursier, 2020-VICO-279314) and the Colin J. Adair Charitable Foundation. HZ is a Wallenberg Scholar supported by grants from the Swedish Research Council (#2022-01018), the European Union’s Horizon Europe research and innovation programme under grant agreement No 101053962, Swedish State Support for Clinical Research (#ALFGBG-71320), the Alzheimer Drug Discovery Foundation (ADDF), USA (#201809-2016862), the AD Strategic Fund and the Alzheimer’s Association (#ADSF-21-831376-C, #ADSF-21-831381-C, and #ADSF-21-831377-C), the Bluefield Project, the Olav Thon Foundation, the Erling-Persson Family Foundation, Stiftelsen för Gamla Tjänarinnor, Hjärnfonden, Sweden (#FO2022-0270), the European Union’s Horizon 2020 research and innovation programme under the Marie Skłodowska-Curie grant agreement No 860197 (MIRIADE), the European Union Joint Programme – Neurodegenerative Disease Research (JPND2021-00694), and the UK Dementia Research Institute at UCL (UKDRI-1003). KB is supported by the Swedish Research Council (#2017-00915 and #2022-00732), the Alzheimer Drug Discovery Foundation (ADDF), USA (#RDAPB-201809-2016615), the Swedish Alzheimer Foundation (#AF-930351, #AF-939721 and #AF-968270), Hjärnfonden, Sweden (#FO2017-0243 and #ALZ2022-0006), the Swedish state under the agreement between the Swedish government and the County Councils, the ALF-agreement (#ALFGBG-715986 and #ALFGBG-965240), the European Union Joint Program for Neurodegenerative Disorders (JPND2019-466-236), the National Institute of Health (NIH), USA, (grant #1R01AG068398-01), the Alzheimer’s Association 2021 Zenith Award (ZEN-21-848495), and the Alzheimer’s Association 2022-2025 Grant (SG-23-1038904 QC). TKK was funded by the Swedish Research Council (Vetenskapsrådet #2021-03244), the Alzheimer’s Association Research Fellowship (#AARF-21-850325), the Swedish Alzheimer Foundation (Alzheimerfonden), the Aina (Ann) Wallströms and Mary-Ann Sjöbloms stiftelsen, and the Emil och Wera Cornells stiftelsen.

## Author contributions

**Lazaro M. Sanchez-Rodriguez:** Conceptualization, Methodology, Software, Formal analysis, Investigation, Data curation, Writing-original draft, Visualization.

**Gleb Bezgin:** Data curation, Formal analysis, Resources, Writing - review & editing.

**Felix Carbonell:** Formal analysis, Resources, Writing - review & editing.

**Joseph Therriault:** Formal analysis, Resources, Writing - review & editing.

**Jaime Fernandez-Arias:** Formal analysis, Resources, Writing - review & editing.

**Stijn Servaes:** Data curation, Writing - review & editing.

**Nesrine Rahmouni:** Data curation, Writing - review & editing.

**Cecile Tissot:** Data curation, Writing - review & editing.

**Jenna Stevenson:** Data curation, Writing - review & editing.

**Thomas K. Karikari:** Data curation, Writing - review & editing.

**Nicholas J. Ashton:** Data curation, Writing - review & editing.

**Andréa L. Benedet:** Data curation, Writing - review & editing.

**Henrik Zetterberg:** Data curation, Writing - review & editing.

**Kaj Blennow:** Data curation, Writing - review & editing.

**Gallen Triana-Baltzer:** Data curation, Writing - review & editing.

**Hartmuth C. Kolb:** Data curation, Writing - review & editing.

**Pedro Rosa-Neto:** Data curation, Resources, Writing - review & editing, Project administration, Funding acquisition.

**Yasser Iturria-Medina:** Conceptualization, Methodology, Investigation, Resources, Writing - review & editing, Supervision, Project administration, Funding acquisition.

## Competing interests

HZ has served at scientific advisory boards and/or as a consultant for Abbvie, Acumen, Alector, Alzinova, ALZPath, Annexon, Apellis, Artery Therapeutics, AZTherapies, CogRx, Denali, Eisai, Nervgen, Novo Nordisk, Optoceutics, Passage Bio, Pinteon Therapeutics, Prothena, Red Abbey Labs, reMYND, Roche, Samumed, Siemens Healthineers, Triplet Therapeutics, and Wave, has given lectures in symposia sponsored by Cellectricon, Fujirebio, Alzecure, Biogen, and Roche, and is a co-founder of Brain Biomarker Solutions in Gothenburg AB (BBS), which is a part of the GU Ventures Incubator Program (outside submitted work). KB has served as a consultant, at advisory boards, or at data monitoring committees for Acumen, ALZPath, BioArctic, Biogen, Eisai, Julius Clinical, Lilly, Novartis, Ono Pharma, Prothena, Roche Diagnostics, and Siemens Healthineers, and is a co-founder of Brain Biomarker Solutions in Gothenburg AB (BBS), which is a part of the GU Ventures Incubator Program, outside the work presented in this paper. The other authors declare no competing interests.

## List of Supplementary Files

Supplementary file 1

Supplementary Figures 1-7, Supplementary Tables 1-7 and Supplementary Pseudocode.

---

**Figure 3—figure supplement 1.**
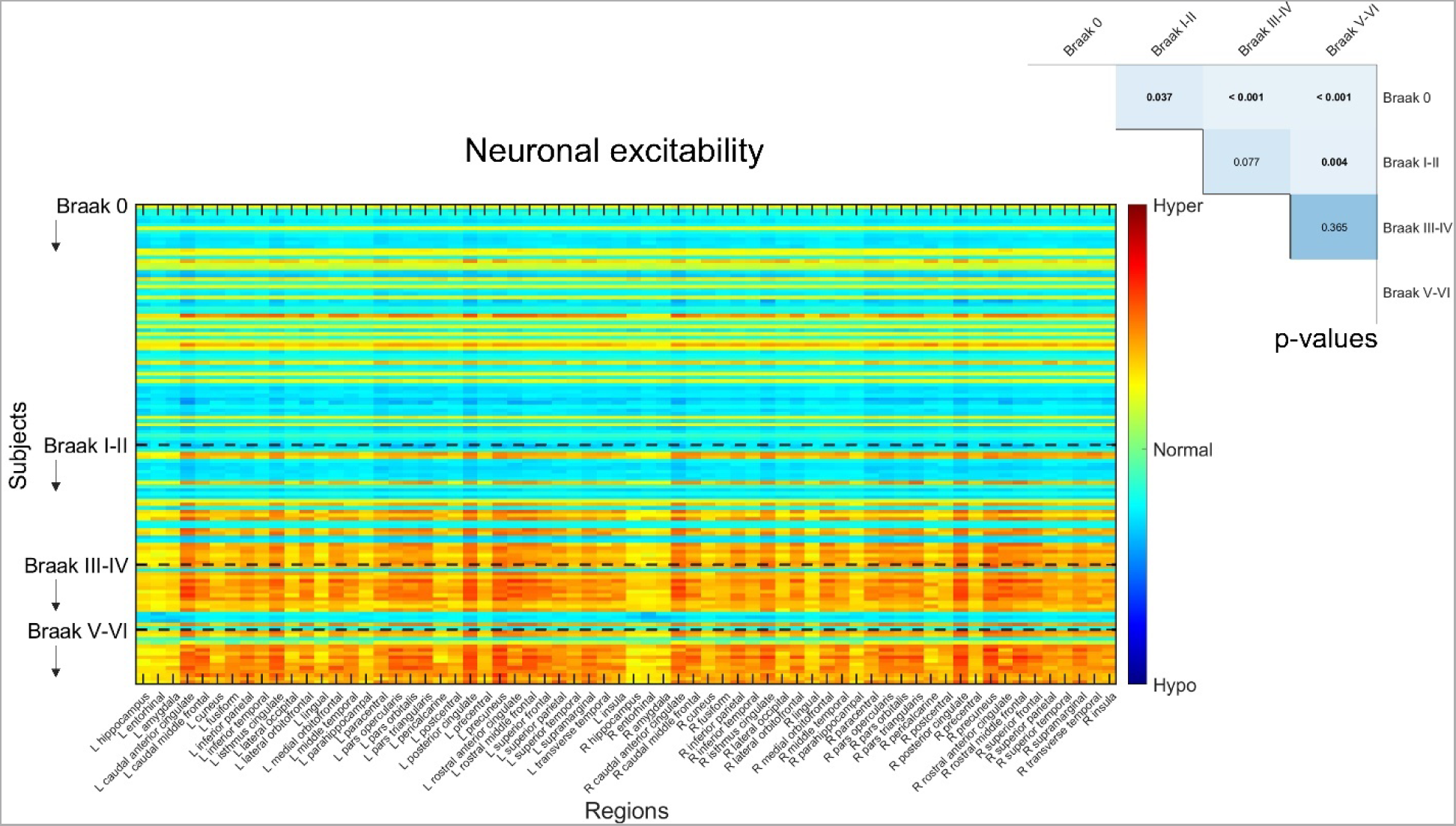
Neuronal excitabilities under the influence of Aβ, tau and Aβ·tau. Participants were grouped according to Braak stages in this figure, to understand tau’s contribution to the individually estimated biological profiles. Inferred neuronal excitability values for the brain regions of interest (“y”-axis) and all subjects (“x”-axis). Within a group, subjects appear according to their existing ordering in the anonymized database. Warm colors represent hyperexcitability of the region in the subject’s brain and cool colors denote hypoexcitable states. Results of ANCOVA post-hoc t-tests for the above-mentioned groups, with the average intra-brain excitability values as response variable and age and sex as covariates appear in the upper right corner. P-values in bold fonts represent differences at a 5% significance level or lower.

## Notes

### Summary of Updates

Introduction, Discussion and Methods updated with further clarifying elements; Figure 2 revised; author affiliations updated; Supplemental files updated.

## References

Abeysuriya, R. G., Hadida, J., Sotiropoulos, S. N., Jbabdi, S., Becker, R., Hunt, B. A. E., Brookes, M. J., & Woolrich, W. (2018). A biophysical model of dynamic balancing of excitation and inhibition in fast oscillatory large-scale networks.

Adewale, Q., Khan, A. F., Carbonell, F., & Iturria-Medina, Y. (2021). Integrated transcriptomic and neuroimaging brain model decodes biological mechanisms in aging and Alzheimer’s disease. ELife, 10. https://doi.org/10.7554/eLife.62589

Archila-Meléndez, M. E., Sorg, C., & Preibisch, C. (2020). Modeling the impact of neurovascular coupling impairments on BOLD-based functional connectivity at rest. NeuroImage, 218(January). https://doi.org/10.1016/j.neuroimage.2020.116871

Ashburner, J. (2007). A fast diffeomorphic image registration algorithm. NeuroImage, 38(1), 95–113. https://doi.org/10.1016/j.neuroimage.2007.07.007

Ashton, N. J., Janelidze, S., Mattsson-Carlgren, N., Binette, A. P., Strandberg, O., Brum, W. S., Karikari, T. K., González-Ortiz, F., di Molfetta, G., Meda, F. J., Jonaitis, E. M., Koscik, R. L., Cody, K., Betthauser, T. J., Li, Y., Vanmechelen, E., Palmqvist, S., Stomrud, E., Bateman, R. J., … Hansson, O. (2022). Differential roles of Aβ42/40, p-tau231 and p-tau217 for Alzheimer’s trial selection and disease monitoring. Nature Medicine. https://doi.org/10.1038/s41591-022-02074-w

Ashton, N. J., Pascoal, T. A., Karikari, T. K., Benedet, A. L., Lantero-Rodriguez, J., Brinkmalm, G., Snellman, A., Schöll, M., Troakes, C., Hye, A., Gauthier, S., Vanmechelen, E., Zetterberg, H., Rosa-Neto, P., & Blennow, K. (2021a). Plasma p-tau231: a new biomarker for incipient Alzheimer’s disease pathology. Acta Neuropathologica, 141(5), 709–724. https://doi.org/10.1007/s00401-021-02275-6

Ashton, N. J., Pascoal, T. A., Karikari, T. K., Benedet, A. L., Lantero-Rodriguez, J., Brinkmalm, G., Snellman, A., Schöll, M., Troakes, C., Hye, A., Gauthier, S., Vanmechelen, E., Zetterberg, H., Rosa-Neto, P., & Blennow, K. (2021b). Plasma p-tau231: a new biomarker for incipient Alzheimer’s disease pathology. Acta Neuropathologica, 141(5), 709–724. https://doi.org/10.1007/s00401-021-<otherinfo> 02275-6</otherinfo>

Babiloni, C., Lizio, R., Del Percio, C., Marzano, N., Soricelli, A., Salvatore, E., Ferri, R., Cosentino, F. I. I., Tedeschi, G., Montella, P., Marino, S., De Salvo, S., Rodriguez, G., Nobili, F., Vernieri, F., Ursini, F., Mundi, C., Richardson, J. C., Frisoni, G. B., & Rossini, P. M. (2013). Cortical Sources of Resting State EEG Rhythms are Sensitive to the Progression of Early Stage Alzheimer’s Disease. Journal of Alzheimer’s Disease, 34(4), 1015–1035. https://doi.org/10.3233/JAD-121750

Benedet, A. L., Milà-Alomà, M., Vrillon, A., Ashton, N. J., Pascoal, T. A., Lussier, F., Karikari, T. K., Hourregue, C., Cognat, E., Dumurgier, J., Stevenson, J., Rahmouni, N., Pallen, V., Poltronetti, N. M., Salvadó, G., Shekari, M., Operto, G., Gispert, J. D., Minguillon, C., … Suárez-Calvet, M. (2021). Differences between Plasma and Cerebrospinal Fluid Glial Fibrillary Acidic Protein Levels across the Alzheimer Disease Continuum. JAMA Neurology, 78(12), 1471–1483. https://doi.org/10.1001/jamaneurol.2021.3671

Bero, A. W., Yan, P., Roh, J. H., Cirrito, J. R., Stewart, F. R., Raichle, M. E., Lee, J. M., & Holtzman, D. M. (2011). Neuronal activity regulates the regional vulnerability to amyloid-β 2 deposition. Nature Neuroscience, 14(6), 750–756. https://doi.org/10.1038/nn.2801

Braak, H., & Braak, E. (1991). Neuropathological stageing of Alzheimer-related changes. Acta Neuropathologica, 82(4), 239–259. https://doi.org/10.1007/BF00308809

Braak, H., Braak, E., & Braak, E. (1995). Staging of Alzheimer’s Disease-Related Neurofibrillary Changes. In Neurobiology of Aging (Vol. 16, Issue 95).

Busche, M. A., & Hyman, B. T. (2020). Synergy between amyloid-β and tau in Alzheimer’s disease. Nature Neuroscience, 23(10), 1183–1193. https://doi.org/10.1038/s41593-020-0687-6

Buxton, R. B., Wong, E. C., & Frank, L. R. (1998). Dynamics of blood flow and oxygenation changes during brain activation: The balloon model. Magnetic Resonance in Medicine, 39(6), 855–864. https://doi.org/10.1002/mrm.1910390602

Celone, K. A., Calhoun, V. D., Dickerson, B. C., Atri, A., Chua, E. F., Miller, S. L., DePeau, K., Rentz, D. M., Selkoe, D. J., Blacker, D., Albert, M. S., & Sperling, R. A. (2006). Alterations in memory networks in mild cognitive impairment and Alzheimer’s disease: An independent component analysis. Journal of Neuroscience, 26(40), 10222–10231. https://doi.org/10.1523/JNEUROSCI.2250-06.2006

Daffertshofer, A., & van Wijk, B. C. M. (2011). On the Influence of Amplitude on the Connectivity between Phases. Frontiers in Neuroinformatics, 5(July), 6. https://doi.org/10.3389/fninf.2011.00006

de Haan, W., van Straaten, E. C. W., Gouw, A. A., & Stam, C. J. (2017). Altering neuronal excitability to preserve network connectivity in a computational model of Alzheimer’s disease. PLoS Computational Biology, 13(9). https://doi.org/10.1371/journal.pcbi.1005707

Deco, G., Cruzat, J., Cabral, J., Knudsen, G. M., Carhart-Harris, R. L., Whybrow, P. C., Logothetis, N. K., & Kringelbach, M. L. (2018). Whole-Brain Multimodal Neuroimaging Model Using Serotonin Receptor Maps Explains Non-linear Functional Effects of LSD. Current Biology, 28(19), 3065–3074.e6. https://doi.org/10.1016/j.cub.2018.07.083

Deco, G., Jirsa, V., Mcintosh, A. R., Sporns, O., & Ko, R. (2009). Key role of coupling, delay, and noise in resting brain fluctuations. 106(25). https://doi.org/10.1073/pnas.0901831106

Deco, G., Kringelbach, M. L., Arnatkeviciute, A., Oldham, S., Sabaroedin, K., Rogasch, N. C., Aquino, K. M., & Fornito, A. (2021). Dynamical consequences of regional heterogeneity in the brain’s transcriptional landscape. In Sci. Adv (Vol. 7). https://www.science.org

Evans, A. C., Kamber, M., Collins, D. L., & MacDonald, D. (1994). An MRI-Based Probabilistic Atlas of Neuroanatomy. In Magnetic Resonance Scanning and Epilepsy (pp. 263–274). Springer US. https://doi.org/10.1007/978-1-4615-2546-2_48

Falcon, M. I., Jirsa, V., & Solodkin, A. (2016). A new neuroinformatics approach to personalized medicine in neurology: The Virtual Brain. In Current Opinion in Neurology (Vol. 29, Issue 4, pp. 429–436). Lippincott Williams and Wilkins. https://doi.org/10.1097/WCO.0000000000000344

Fernández Arias, J., Therriault, J., Thomas, E., Lussier, F. Z., Bezgin, G., Tissot, C., Servaes, S., Mathotaarachchi, S. S., Schoemaker, D., Stevenson, J., Rahmouni, N., Kang, M. S., Pallen, V., Poltronetti, N. M., Wang, Y.-T., Kunach, P., Chamoun, M., Quispialaya S, K. M., Vitali, P., … Rosa-Neto, P. (2023). Verbal memory formation across PET-based Braak stages of tau accumulation in Alzheimer’s disease. Brain Communications, 5(3). https://doi.org/10.1093/braincomms/fcad146

Folstein, M. F., Folstein, S. E., & Mchugh, P. R. (1975). ‘MINI-MENTAL STATE’ A PRACTICAL METHOD FOR GRADING THE COGNITIVE STATE OF PATIENTS FOR THE CLINICIAN*. In J. gsychiaf. Res (Vol. 12). Pergamon Press.

Friston, K. J., Mechelli, A., Turner, R., & Price, C. J. (2000). Nonlinear responses in fMRI: The balloon model, Volterra kernels, and other hemodynamics. NeuroImage, 12(4), 466–477. https://doi.org/10.1006/nimg.2000.0630

Gjorgjieva, J., Evers, J. F., & Eglen, S. J. (2016). Homeostatic activity-dependent tuning of recurrent networks for robust propagation of activity. Journal of Neuroscience, 36(13), 3722–3734. https://doi.org/10.1523/JNEUROSCI.2511-15.2016

Insel, P. S., Mormino, E. C., Aisen, P. S., Thompson, W. K., & Donohue, M. C. (2020). Neuroanatomical spread of amyloid β and tau in Alzheimer’s disease: implications for primary prevention. Brain Communications, 2(1), 1–11. https://doi.org/10.1093/braincomms/fcaa007

Iturria-Medina, Y., Canales-Rodríguez, E. J., Melie-García, L., Valdés-Hernández, P. A., Martínez-Montes, E., Alemán-Gómez, Y., & Sánchez-Bornot, J. M. (2007). Characterizing brain anatomical connections using diffusion weighted MRI and graph theory. NeuroImage, 36(3), 645–660. https://doi.org/10.1016/j.neuroimage.2007.02.012

Iturria-Medina, Y., Carbonell, F., Assadi, A., Adewale, Q., Khan, A. F., Baumeister, T. R., & Sanchez-Rodriguez, L. (2021). Integrating molecular, histopathological, neuroimaging and clinical neuroscience data with NeuroPM-box. Communications Biology, 4(1). https://doi.org/10.1038/s42003-021-02133-x

Iturria-Medina, Y., Carbonell, F. M., & Evans, A. C. (2018). Multimodal imaging-based therapeutic fingerprints for optimizing personalized interventions: Application to neurodegeneration. NeuroImage, 179(May), 40–50. https://doi.org/10.1016/j.neuroimage.2018.06.028

Iturria-Medina, Y., Carbonell, F. M., Sotero, R. C., Chouinard-Decorte, F., & Evans, A. C. (2017). Multifactorial causal model of brain (dis)organization and therapeutic intervention: Application to Alzheimer’s disease. NeuroImage, 152(February), 60–77. https://doi.org/10.1016/j.neuroimage.2017.02.058

Iturria-Medina, Y., & Evans, A. C. (2015). On the central role of brain connectivity in neurodegenerative disease progression. In Frontiers in Aging Neuroscience (Vol. 7, Issue MAY). Frontiers Media S.A. https://doi.org/10.3389/fnagi.2015.00090

Iturria-Medina, Y., & Evans, A. C. (2021). Networks-Mediated Spreading of Pathology in Neurodegenerative Diseases. In Brain Network Dysfunction in Neuropsychiatric Illness (pp. 171– 186). Springer International Publishing. https://doi.org/10.1007/978-3-030-59797-9_9

Iturria-Medina, Y., Sotero, R. C., Toussaint, P. J., Mateos-Perez, J. M., Evans, A. C., & Initiative, T. A. D. N. (2016). Early role of vascular dysregulation on late-onset Alzheimer’s disease based on multifactorial data-driven analysis. Nat Commun, 7(May), 11934. https://doi.org/10.1038/ncomms11934

Jack, C. R., Bennett, D. A., Blennow, K., Carrillo, M. C., Dunn, B., Haeberlein, S. B., Holtzman, D. M., Jagust, W., Jessen, F., Karlawish, J., Liu, E., Molinuevo, J. L., Montine, T., Phelps, C., Rankin, K. P., Rowe, C. C., Scheltens, P., Siemers, E., Snyder, H. M., … Silverberg, N. (2018). NIA-AA Research Framework: Toward a biological definition of Alzheimer’s disease. In Alzheimer’s and Dementia (Vol. 14, Issue 4, pp. 535–562). Elsevier Inc. https://doi.org/10.1016/j.jalz.2018.02.018

Jagust, W. J., Bandy, D., Chen, K., Foster, N. L., Landau, S. M., Mathis, C. A., Price, J. C., Reiman, E. M., Skovronsky, D., & Koeppe, R. A. (2010). The Alzheimer’s Disease Neuroimaging Initiative positron emission tomography core. Alzheimer’s and Dementia, 6(3), 221–229. https://doi.org/10.1016/j.jalz.2010.03.003

Jansen, B. H., & Rit, V. G. (1995). Electroencephalogram and visual evoked potential generation in a mathematical model of coupled cortical columns. Biological Cybernetics, 73(4), 357–366. https://doi.org/10.1007/BF00199471

Jia, X. Z., Wang, J., Sun, H. Y., Zhang, H., Liao, W., Wang, Z., Yan, C. G., Song, X. W., & Zang, Y. F. (2019). RESTplus: an improved toolkit for resting-state functional magnetic resonance imaging data processing. In Science Bulletin (Vol. 64, Issue 14, pp. 953–954). Elsevier B.V. https://doi.org/10.1016/j.scib.2019.05.008

Karikari, T. K., Ashton, N. J., Rodriguez, J. L., Schöll, M., Höglund, K., Brinkmalm, G., Zetterberg, H., Blennow, K., A Pascoal, C. T., Benedet, A. L., Chamoun, M., Savard, M., Kang, M. S., Therriault, J., Gauthier, S., Rosa-Neto, P., Pascoal, T. A., Masserweh, G., Soucy, J., … Blennow, K. (2020). Blood phosphorylated tau 181 as a biomarker for Alzheimer’s disease: a diagnostic performance and prediction modelling study using data from four prospective cohorts. In Articles Lancet Neurol (Vol. 19). www.thelancet.com/neurology

Kazim, S. F., Chuang, S. C., Zhao, W., Wong, R. K. S., Bianchi, R., & Iqbal, K. (2017). Early-onset network hyperexcitability in presymptomatic Alzheimer’s disease transgenic mice is suppressed by passive immunization with anti-human APP/Aβ antibody and by mGluR5 blockade. Frontiers in Aging Neuroscience, 9(MAR). https://doi.org/10.3389/fnagi.2017.00071

Khan, A. F., Adewale, Q., Baumeister, T. R., Carbonell, F., Zilles, K., Palomero-Gallagher, N., & Iturria-Medina, Y. (2022). Personalized brain models identify neurotransmitter receptor changes in Alzheimer’s disease. Brain: A Journal of Neurology, 145(5), 1785–1804. https://doi.org/10.1093/brain/awab375

Klein, A., & Tourville, J. (2012). 101 Labeled Brain Images and a Consistent Human Cortical Labeling Protocol. Frontiers in Neuroscience, 6(DEC), 1–12. https://doi.org/10.3389/fnins.2012.00171

Kwon, H. S., & Koh, S. H. (2020). Neuroinflammation in neurodegenerative disorders: the roles of microglia and astrocytes. In Translational Neurodegeneration (Vol. 9, Issue 1). BioMed Central Ltd. https://doi.org/10.1186/s40035-020-00221-2

Lauterborn, J. C., Scaduto, P., Cox, C. D., Schulmann, A., Lynch, G., Gall, C. M., Keene, C. D., & Limon, A. (2021). Increased excitatory to inhibitory synaptic ratio in parietal cortex samples from individuals with Alzheimer’s disease. Nature Communications, 12(1), 2603. https://doi.org/10.1038/s41467-021-22742-8

Logothetis, N. K., Pauls, J., Augath, M., Trinath, T., & Oeltermann, A. (2001). Neurophysiological investigation of the basis of the fMRI signal.

Luppi, A. I., Cabral, J., Cofre, R., Destexhe, A., Deco, G., & Kringelbach, M. L. (2022). Dynamical models to evaluate structure–function relationships in network neuroscience. In Nature Reviews Neuroscience. Springer Nature. https://doi.org/10.1038/s41583-022-00646-w

Maestú, F., Cuesta, P., Hasan, O., Fernandéz, A., Funke, M., & Schulz, P. E. (2019). The Importance of the Validation of M/EEG With Current Biomarkers in Alzheimer’s Disease. Frontiers in Human Neuroscience, 13(February), 1–10. https://doi.org/10.3389/fnhum.2019.00017

Maestú, F., de Haan, W., Busche, M. A., & DeFelipe, J. (2021). Neuronal Excitation/Inhibition imbalance: a core element of a translational perspective on Alzheimer pathophysiology. Ageing Research Reviews, 69, 101372. https://doi.org/10.1016/j.arr.2021.101372

Mederos, S., & Perea, G. (2019). GABAergic-astrocyte signaling: A refinement of inhibitory brain networks. In GLIA (Vol. 67, Issue 10, pp. 1842–1851). John Wiley and Sons Inc. https://doi.org/10.1002/glia.23644

Meijer, H. G. E., Eissa, T. L., Kiewiet, B., Neuman, J. F., Schevon, C. A., Emerson, R. G., Goodman, R. R., McKhann, G. M., Marcuccilli, C. J., Tryba, A. K., Cowan, J. D., van Gils, S. A., & van Drongelen, W. (2015). Modeling focal epileptic activity in the Wilson-cowan model with depolarization block. Journal of Mathematical Neuroscience, 5, 7. https://doi.org/10.1186/s13408-015-0019-4

Milà-Alomà, M., Ashton, N. J., Shekari, M., Salvadó, G., Ortiz-Romero, P., Montoliu-Gaya, L., Benedet, A. L., Karikari, T. K., Lantero-Rodriguez, J., Vanmechelen, E., Day, T. A., González-Escalante, A., Sánchez-Benavides, G., Minguillon, C., Fauria, K., Molinuevo, J. L., Dage, J. L., Zetterberg, H., Gispert, J. D., … Blennow, K. (2022). Plasma p-tau231 and p-tau217 as state markers of amyloid-β pathology in preclinical Alzheimer’s disease. Nature Medicine, 28(9), 1797–1801. https://doi.org/10.1038/s41591-022-01925-w

Nasreddine, Z. S., Phillips, N. A., Bédirian, V., Charbonneau, S., Whitehead, V., Collin, I., Cummings, J. L., & Chertkow, H. (2005). The Montreal Cognitive Assessment, MoCA: A brief screening tool for mild cognitive impairment. Journal of the American Geriatrics Society, 53(4), 695–699. https://doi.org/10.1111/j.1532-5415.2005.53221.x

Nichols, K. J., Chen, B., Tomas, M. B., & Palestro, C. J. (2018). Interpreting 123I–ioflupane dopamine transporter scans using hybrid scores. European Journal of Hybrid Imaging, 2(1). https://doi.org/10.1186/s41824-018-0028-0

Nutma, E., Fancy, N., Weinert, M., Marzin, M. C., Muirhead, R. C., Falk, I., de Bruin, J., Hollaus, D., Anink, J., Story, D., Chandran, S., Tang, J., Saito, T., Saido, T. C., Wiltshire, K., Beltran-Lobo, P., Philips, A., Antel, J., Healy, L., … Owen, D. (n.d.-a). Translocator protein is a marker of activated microglia in rodent models but not human neurodegenerative diseases. https://doi.org/10.1101/2022.05.11.491453

Nutma, E., Fancy, N., Weinert, M., Marzin, M. C., Muirhead, R. C., Falk, I., de Bruin, J., Hollaus, D., Anink, J., Story, D., Chandran, S., Tang, J., Saito, T., Saido, T. C., Wiltshire, K., Beltran-Lobo, P., Philips, A., Antel, J., Healy, L., … Owen, D. (n.d.-b). Translocator protein is a marker of activated microglia in rodent models but not human neurodegenerative diseases. https://doi.org/10.1101/2022.05.11.491453

Obata, T., Liu, T. T., Miller, K. L., Luh, W., Wong, E. C., Frank, L. R., & Buxton, R. B. (2004). Discrepancies between BOLD and flow dynamics in primary and supplementary motor areas: application of the balloon model to the interpretation of BOLD transients. 21, 144–153. https://doi.org/10.1016/j.neuroimage.2003.08.040

Ossenkoppele, R., Pichet Binette, A., Groot, C., Smith, R., Strandberg, O., Palmqvist, S., Stomrud, E., Tideman, P., Ohlsson, T., Jögi, J., Johnson, K., Sperling, R., Dore, V., Masters, C. L., Rowe, C., Visser, D., van Berckel, B. N. M., van der Flier, W. M., Baker, S., … Hansson, O. (2022). Amyloid and tau PET-positive cognitively unimpaired individuals are at high risk for future cognitive decline. Nature Medicine, 28(11), 2381–2387. https://doi.org/10.1038/s41591-022-02049-x

Pascoal, T. A., Benedet, A. L., Ashton, N. J., Kang, M. S., Therriault, J., Chamoun, M., Savard, M., Lussier, F. Z., Tissot, C., Karikari, T. K., Ottoy, J., Mathotaarachchi, S., Stevenson, J., Massarweh, G., Schöll, M., de Leon, M. J., Soucy, J. P., Edison, P., Blennow, K., … Rosa-Neto, P. (2021). Microglial activation and tau propagate jointly across Braak stages. Nature Medicine, 27(9), 1592– 1599. https://doi.org/10.1038/s41591-021-01456-w

Pascoal, T. A., Therriault, J., Benedet, A. L., Savard, M., Lussier, F. Z., Chamoun, M., Tissot, C., Qureshi, M. N. I., Kang, M. S., Mathotaarachchi, S., Stevenson, J., Hopewell, R., Massarweh, G., Soucy, J.-P., Gauthier, S., & Rosa-Neto, P. (2020). 18F-MK-6240 PET for early and late detection of neurofibrillary tangles. Brain, 143(9), 2818–2830. https://doi.org/10.1093/brain/awaa180

Picconi, B., Piccoli, G., & Calabresi, P. (2012). Synaptic Dysfunction in Parkinson’s Disease (pp. 553– 572). https://doi.org/10.1007/978-3-7091-0932-8_24

Roshanbin, S., Xiong, M., Hultqvist, G., Söderberg, L., Zachrisson, O., Meier, S., Ekmark-Lewén, S., Bergström, J., Ingelsson, M., Sehlin, D., & Syvänen, S. (2022). In vivo imaging of alpha-synuclein with antibody-based PET. Neuropharmacology, 208. https://doi.org/10.1016/j.neuropharm.2022.108985

Sanchez-Rodriguez, L. M., Iturria-Medina, Y., Baines, E. A., Mallo, S. C., Dousty, M., & Sotero, R. C. (2018). Design of optimal nonlinear network controllers for Alzheimer’s disease. PLOS Computational Biology, 14(5), e1006136. https://doi.org/10.1371/journal.pcbi.1006136

Sanchez-Rodriguez, L. M., Iturria-Medina, Y., Mouches, P., & Sotero, R. C. (2021). Detecting brain network communities: Considering the role of information flow and its different temporal scales. NeuroImage, 225(Jan), 117431. https://doi.org/10.1016/j.neuroimage.2020.117431

Shen, Z., Bao, X., & Wang, R. (2018). Clinical PET imaging of microglial activation: Implications for microglial therapeutics in Alzheimer’s disease. In Frontiers in Aging Neuroscience (Vol. 10, Issue OCT). Frontiers Media S.A. https://doi.org/10.3389/fnagi.2018.00314

Simon, A. B., & Buxton, R. B. (2015). Understanding the dynamic relationship between cerebral blood flow and the BOLD signal: Implications for quantitative functional MRI. NeuroImage, 116, 158–167. https://doi.org/10.1016/j.neuroimage.2015.03.080

Sotero, R. C., & Trujillo-Barreto, N. J. (2007). Modelling the role of excitatory and inhibitory neuronal activity in the generation of the BOLD signal. NeuroImage, 35(1), 149–165. https://doi.org/10.1016/j.neuroimage.2006.10.027

Sotero, R. C., & Trujillo-Barreto, N. J. (2008). Biophysical model for integrating neuronal activity, EEG, fMRI and metabolism. NeuroImage, 39, 290–309. https://doi.org/10.1016/j.neuroimage.2007.08.001

Sotero, R. C., Trujillo-Barreto, N. J., Jiménez, J. C., Carbonell, F., & Rodríguez-Rojas, R. (2009). Identification and comparison of stochastic metabolic/hemodynamic models (sMHM) for the generation of the BOLD signal. Journal of Computational Neuroscience, 26(2), 251–269. https://doi.org/10.1007/s10827-008-0109-3

Stefanovski, L., Triebkorn, P., Spiegler, A., Diaz-Cortes, M. A., Solodkin, A., Jirsa, V., McIntosh, A. R., & Ritter, P. (2019). Linking Molecular Pathways and Large-Scale Computational Modeling to Assess Candidate Disease Mechanisms and Pharmacodynamics in Alzheimer’s Disease. Frontiers in Computational Neuroscience, 13(August), 1–27. https://doi.org/10.3389/fncom.2019.00054

Tan Toi, P., Jae Jang, H., Min, K., Kim, S.-P., Lee, S.-K., Lee, J., Kwag, J., & Park, J.-Y. (n.d.). In vivo direct imaging of neuronal activity at high temporospatial resolution. https://www.science.org

Targa Dias Anastacio, H., Matosin, N., & Ooi, L. (2022). Neuronal hyperexcitability in Alzheimer’s disease: what are the drivers behind this aberrant phenotype? In Translational Psychiatry (Vol. 12, Issue 1). Springer Nature. https://doi.org/10.1038/s41398-022-02024-7

Therriault, J., Benedet, A. L., Pascoal, T. A., Savard, M., Ashton, N. J., Chamoun, M., Tissot, C., Lussier, F., Kang, M. S., Bezgin, G., Wang, T., Fernandes-Arias, J., Massarweh, G., Vitali, P., Zetterberg, H., Blennow, K., Saha-Chaudhuri, P., Soucy, J. P., Gauthier, S., & Rosa-Neto, P. (2021). Determining amyloid-b positivity using 18F-AZD4694 PET imaging. In Journal of Nuclear Medicine (Vol. 62, Issue 2, pp. 247–252). Society of Nuclear Medicine Inc. https://doi.org/10.2967/jnumed.120.245209

Therriault, J., Pascoal, T. A., Lussier, F. Z., Tissot, C., Chamoun, M., Bezgin, G., Servaes, S., Benedet, A. L., Ashton, N. J., Karikari, T. K., Lantero-Rodriguez, J., Kunach, P., Wang, Y. T., Fernandez-Arias, J., Massarweh, G., Vitali, P., Soucy, J. P., Saha-Chaudhuri, P., Blennow, K., … Rosa-Neto, P. (2022). Biomarker modeling of Alzheimer’s disease using PET-based Braak staging. Nature Aging, 2(6), 526–535. https://doi.org/10.1038/s43587-022-00204-0

Therriault, J., Vermeiren, M., Servaes, S., Tissot, C., Ashton, N. J., Benedet, A. L., Karikari, T. K., Lantero-Rodriguez, J., Brum, W. S., Lussier, F. Z., Bezgin, G., Stevenson, J., Rahmouni, N., Kunach, P., Wang, Y.-T., Fernandez-Arias, J., Socualaya, K. Q., Macedo, A. C., Ferrari-Souza, J. P., … Rosa-Neto, P. (2022). Association of Phosphorylated Tau Biomarkers With Amyloid Positron Emission Tomography vs Tau Positron Emission Tomography. JAMA Neurology. https://doi.org/10.1001/jamaneurol.2022.4485

Tissot, C., L. Benedet, A., Therriault, J., Pascoal, T. A., Lussier, F. Z., Saha-Chaudhuri, P., Chamoun, M., Savard, M., Mathotaarachchi, S. S., Bezgin, G., Wang, Y. T., Fernandez Arias, J., Rodriguez, J. L., Snellman, A., Ashton, N. J., Karikari, T. K., Blennow, K., Zetterberg, H., de Villers-Sidani, E., … Rosa-Neto, P. (2021). Plasma pTau181 predicts cortical brain atrophy in aging and Alzheimer’s disease. Alzheimer’s Research and Therapy, 13(1). https://doi.org/10.1186/s13195-021-00802-x

Tissot, C., Servaes, S., Lussier, F., Pedro Ferrari Souza, J., Therriault, J., Cristina Lukasewicz Ferreira, P., Bezgin, G., Bellaver, B., Teixeira Leffa, D., Mathotaarachchi, S. S., Stevenson, J. B., Rahmouni, N., Su Kang, M., Pallen, V. B., Margherita-Poltronetti, N., Wang, Y.-T., Fernandez-Arias, J., Benedet, A. L., Zimmer, E.R., … Professor of Psychiatry, A. (n.d.). The association of age-related and off-target retention with longitudinal quantification of [18 F]MK6240 tau-PET in target regions. https://doi.org/10.1101/2022.05.24.22275386

Tok, S., Maurin, H., Delay, C., Crauwels, D., Manyakov, N. V., Van Der Elst, W., Moechars, D., & Drinkenburg, W. H. I. M. (2022). Pathological and neurophysiological outcomes of seeding human-derived tau pathology in the APP-KI NL-G-F and NL-NL mouse models of Alzheimer’s Disease. Acta Neuropathologica Communications, 10(1). https://doi.org/10.1186/s40478-022-01393-w

Tournier, J. D., Yeh, C. H., Calamante, F., Cho, K. H., Connelly, A., & Lin, C. P. (2008). Resolving crossing fibres using constrained spherical deconvolution: Validation using diffusion-weighted imaging phantom data. NeuroImage, 42(2), 617–625. https://doi.org/10.1016/j.neuroimage.2008.05.002

Triana-Baltzer, G., Moughadam, S., Slemmon, R., van Kolen, K., Theunis, C., Mercken, M., & Kolb, H. C. (2021). Development and validation of a high-sensitivity assay for measuring p217+tau in plasma. *Alzheimer’s and Dementia: Diagnosis*, Assessment and Disease Monitoring, 13(1). https://doi.org/10.1002/dad2.12204

Valdes-Sosa, P. A., Sanchez-Bornot, J. M., Sotero, R. C., Iturria-Medina, Y., Aleman-Gomez, Y., Bosch-Bayard, J., Carbonell, F., & Ozaki, T. (2009). Model driven EEG/fMRI fusion of brain oscillations. Human Brain Mapping, 30(9), 2701–2721. https://doi.org/10.1002/hbm.20704

van Nifterick, A. M., Gouw, A. A., van Kesteren, R. E., Scheltens, P., Stam, C. J., & de Haan, W. (2022). A multiscale brain network model links Alzheimer’s disease-mediated neuronal hyperactivity to large-scale oscillatory slowing. Alzheimer’s Research & Therapy, 14(1), 101. https://doi.org/10.1186/s13195-022-01041-4

Vogel, J. W., Iturria-Medina, Y., Strandberg, O. T., Smith, R., Levitis, E., Evans, A. C., & Hansson, O. (2020). Spread of pathological tau proteins through communicating neurons in human Alzheimer’s disease. Nature Communications, 11(1), 2612. https://doi.org/10.1038/s41467-020-15701-2

Vossel, K. A., Tartaglia, M. C., Nygaard, H. B., Zeman, A. Z., & Miller, B. L. (2017). Epileptic activity in Alzheimer’s disease: causes and clinical relevance. The Lancet Neurology, 16(4), 311–322. https://doi.org/10.1016/S1474-4422(17)30044-3

Wilson, H. R., & Cowan, J. D. (1972). Excitatory and inhibitory interactions in localized populations of model neurons. Biophysical Journal, 12(1), 1–24. https://doi.org/10.1016/S0006-3495(72)86068-5

Yang, L., Yan, Y., Li, Y., Hu, X., Lu, J., Chan, P., Yan, T., & Han, Y. (2020). Frequency-dependent changes in fractional amplitude of low-frequency oscillations in Alzheimer’s disease: a resting-state fMRI study. Brain Imaging and Behavior, 14(6), 2187–2201. https://doi.org/10.1007/s11682-019-00169-6

Yang, L., Yan, Y., Wang, Y., Hu, X., Lu, J., Chan, P., Yan, T., & Han, Y. (2018). Gradual Disturbances of the Amplitude of Low-Frequency Fluctuations (ALFF) and Fractional ALFF in Alzheimer Spectrum. Frontiers in Neuroscience, 12(December), 1–16. https://doi.org/10.3389/fnins.2018.00975

Young, P. N. E., Estarellas, M., Coomans, E., Srikrishna, M., Beaumont, H., Maass, A., Venkataraman, A. v., Lissaman, R., Jiménez, D., Betts, M. J., McGlinchey, E., Berron, D., O’Connor, A., Fox, N. C., Pereira, J. B., Jagust, W., Carter, S. F., Paterson, R. W., & Schöll, M. (2020). Imaging biomarkers in neurodegeneration: Current and future practices. In Alzheimer’s Research and Therapy (Vol. 12, Issue 1). BioMed Central Ltd. https://doi.org/10.1186/s13195-020-00612-7

Zimmermann, J., Perry, A., Breakspear, M., Schirner, M., Sachdev, P., Wen, W., Kochan, N. A., Mapstone, M., Ritter, P., McIntosh, A. R., & Solodkin, A. (2018). Differentiation of Alzheimer’s disease based on local and global parameters in personalized Virtual Brain models. NeuroImage: Clinical, 19, 240–251. https://doi.org/10.1016/j.nicl.2018.04.017

