## Supplementary Information for "Revealing the combined roles of Aβ and tau in Alzheimer’s disease via a pathophysiological activity decoder"

Sanchez-Rodriguez et al.

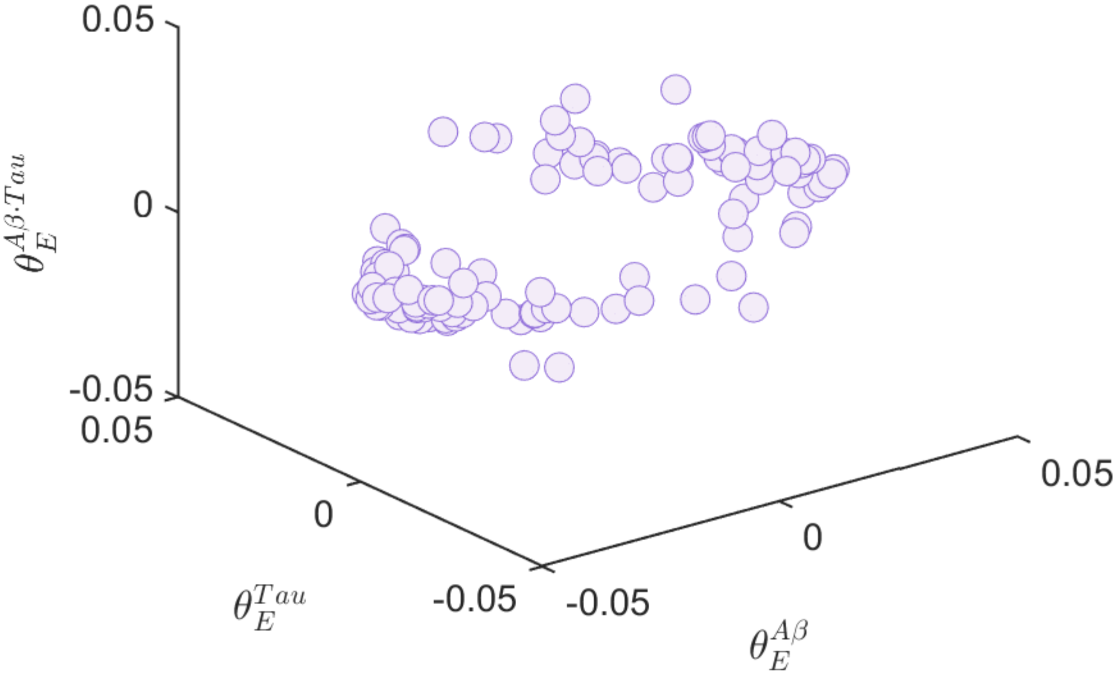


**Supplementary file 1—figure 1. Estimated neuronal activity influences by *Aβ, tau and Aβ·tau*.** Each dot represents a subject-specific set of pathophysiological effects on neuronal excitability, obtained through parameter estimation following the model of equation (1) in the main text. The axes correspond to the Aβ, tau and Aβ·tau weights, respectively.

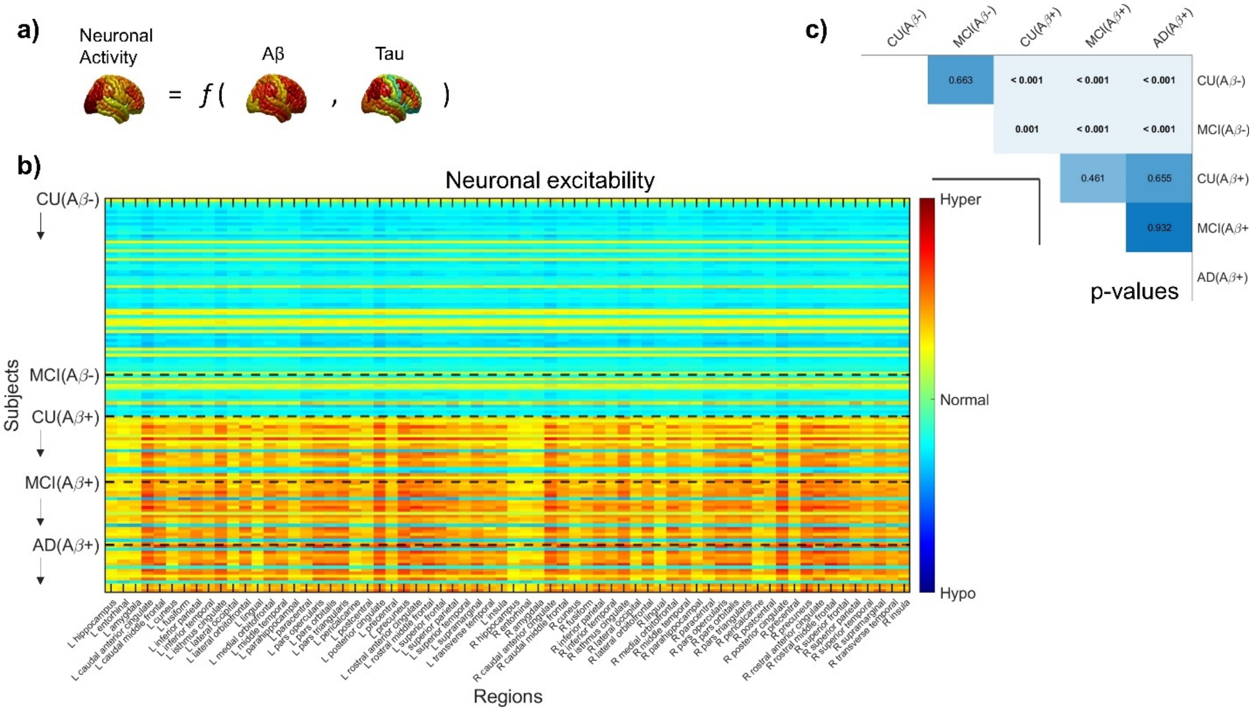


**Supplementary file 1—figure 2. Neuronal excitabilities under the influence of *Aβ and tau* (participants grouped according to Aβ-positivity). a**) Schematic representation of the influence model. **b**) Inferred neuronal excitability values for the brain regions of interest (“y”-axis) and all subjects (“x”-axis). Within a group, subjects appear according to their existing ordering in the anonymized database. Warm colors represent hyperexcitability of the region in the subject’s brain and cool colors denote hypoexcitable states. **c**) Results of ANCOVA post-hoc t-tests for the above-mentioned groups, with the average intra-brain excitability values as response variable and age and sex as covariates. P-values in bold fonts represent differences at a 5% significance level or lower.

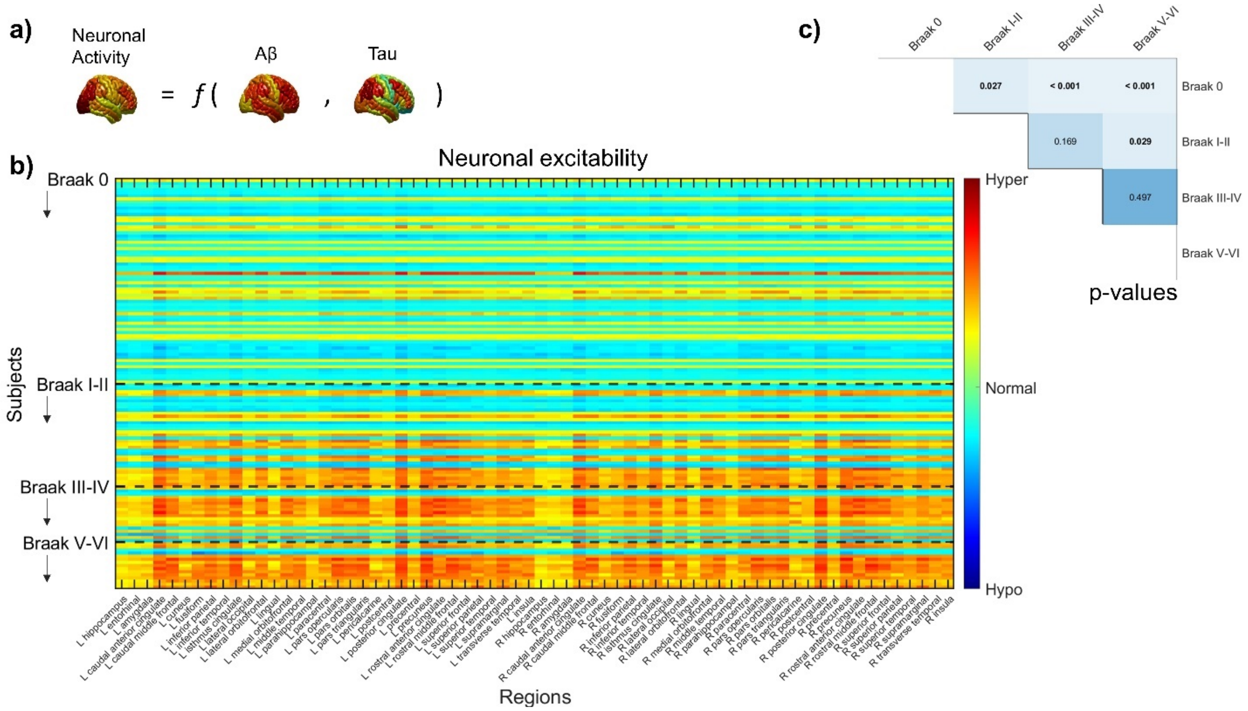


**Supplementary file 1—figure 3. Neuronal excitabilities under the influence of *Aβ and tau* (participants grouped according to Braak stages). a**) Schematic representation of the influence model. **b**) Inferred neuronal excitability values for the brain regions of interest ("y"-axis) and all subjects ("x"-axis). Within a group, subjects appear according to their existing ordering in the anonymized database. Warm colors represent hyperexcitability of the region in the subject's brain and cool colors denote hypoexcitable states. **c**) Results of ANCOVA post-hoc t-tests for the above-mentioned groups, with the average intra-brain excitability values as response variable and age and sex as covariates. P-values in bold fonts represent differences at a 5% significance level or lower.

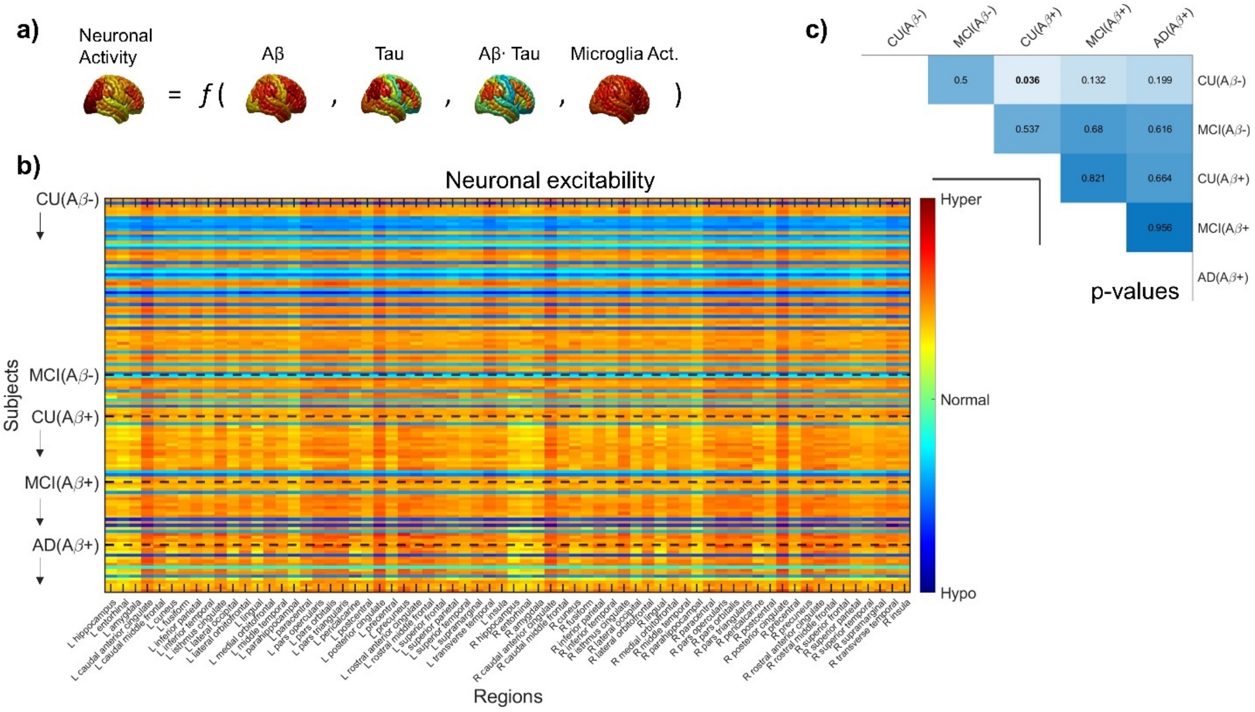


**Supplementary file 1—figure 4. Neuronal excitabilities under the influence of *Aβ, tau, Aβ·tau and microglial activation* (participants grouped according to Aβ-positivity). a**) Schematic representation of the influence model. **b**) Inferred neuronal excitability values for the brain regions of interest ("y"-axis) and all subjects ("x"-axis). Within a group, subjects appear according to their existing ordering in the anonymized database. Warm colors represent hyperexcitability of the region in the subject's brain and cool colors denote hypoexcitable states. **c**) Results of ANCOVA post-hoc t-tests for the above-mentioned groups, with the average intra-brain excitability values as response variable and age and sex as covariates. No significant differences between groups were observed.

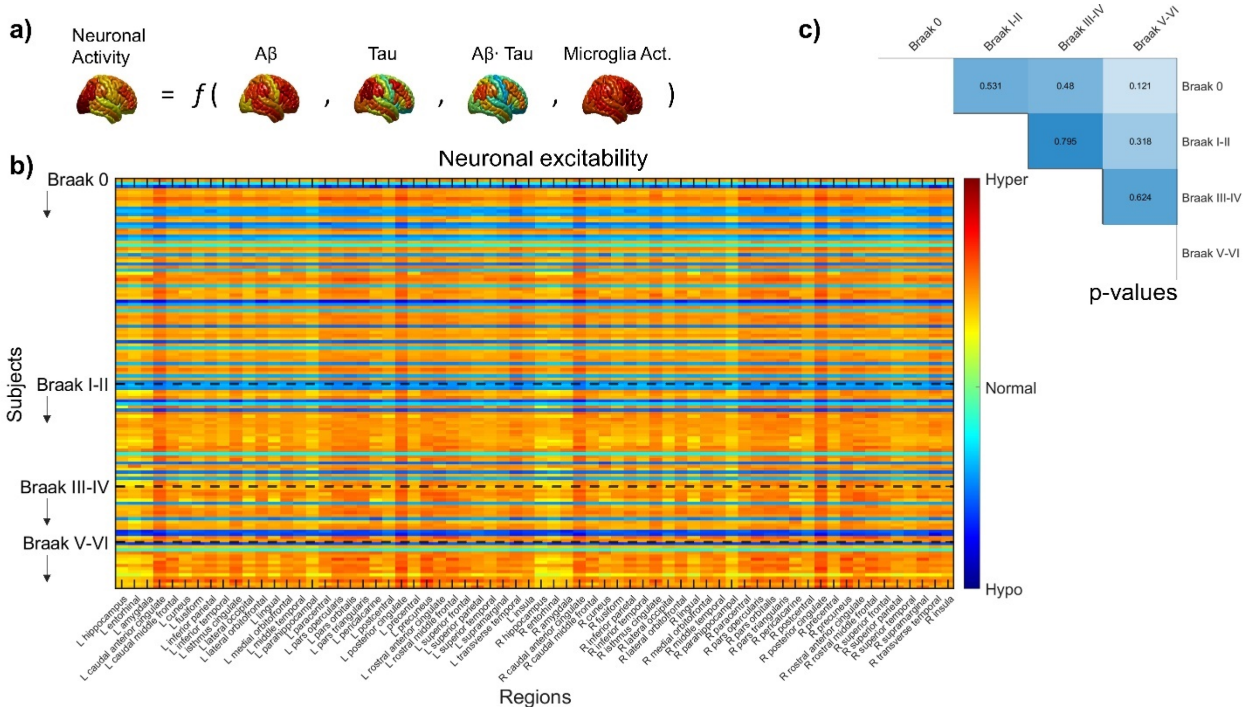


**Supplementary file 1—figure 5. Neuronal excitabilities under the influence of *Aβ, tau, Aβ·tau and microglial activation* (participants grouped according to Braak stages). a**) Schematic representation of the influence model. **b**) Inferred neuronal excitability values for the brain regions of interest (“y”-axis) and all subjects (“x”-axis). Within a group, subjects appear according to their existing ordering in the anonymized database. Warm colors represent hyperexcitability of the region in the subject’s brain and cool colors denote hypoexcitable states. **c**) Results of ANCOVA post-hoc t-tests for the above-mentioned groups, with the average intra-brain excitability values as response variable and age and sex as covariates. P-values in bold fonts represent differences at a 5% significance level or lower.

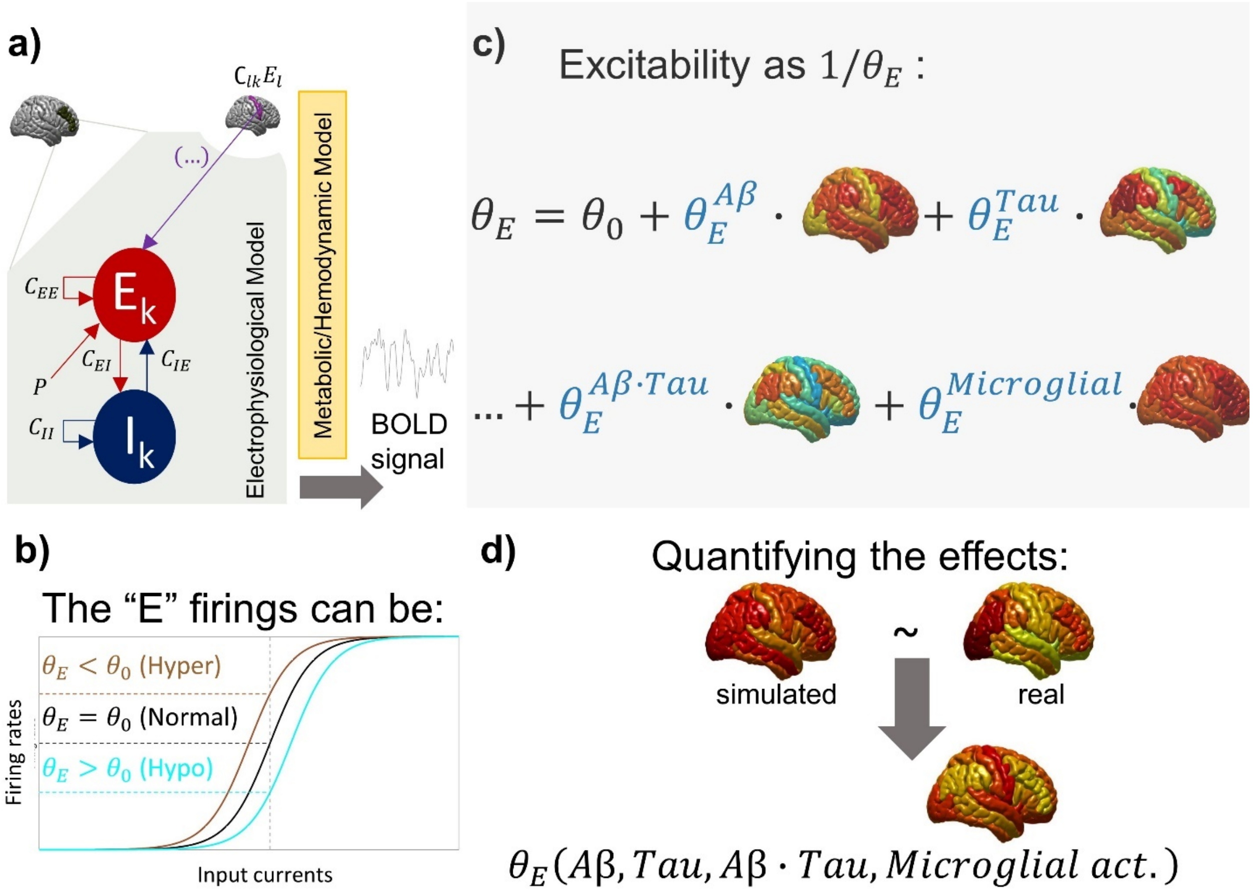


**Supplementary file 1—figure 6. Overview of integrating pathophysiological information into the computational brain activity decoder for Alzheimer's. a**) Each brain region, $k$, is modeled as coupled excitatory and inhibitory populations. For example, $C_{EI}$ represents the strength of the excitatory connection with the inhibitory neural mass. Additionally, the excitatory population receives an unspecified local stimulus that accounts for unmodeled interactions, $P$, and excitatory inputs from other regions via a connectivity matrix obtained from diffusion MRI ($C_{lk}$). The excitatory and inhibitory inputs feed a metabolic/hemodynamic model that simulates the resting-state BOLD signal (fMRI) for the given region. **b**) In the study, the excitatory firing rates are obtained by using sigmoid functions with varying firing thresholds. A low sigmoidal threshold value means high firing rates ("hyperexcitable state") at a given input current, while the opposite is termed "hypoexcitability", as compared to baseline firing conditions. **c**) The subject-specific influence of pathophysiological factors on neuronal activity is modeled as linear changes on a region's firing threshold due to the competition/contribution of the regional pathological loads. In Alzheimer's disease, we assume that the threshold can be, in general, affected by Aβ plaques, tau tangles, the interaction of Aβ and tau (modeled as the regional multiplication of the participant's Aβ and tau SUVRs), and microglial activation. Thus, the anatomical representations in the figure gather PET SUVRs corresponding to these factors in 66 regions of interest. The proportionality constants ($\theta_E^{Ab}$, $\theta_E^{Tau}$, $\theta_E^{Ab \cdot Tau}$, $\theta_E^{Microglial\ act.}$) characterize the subject-specific global influence of the pathologies. **d**) We infer these pathological influences by minimizing the similarity between the observed neuronal activity biomarkers (i.e., the fractional amplitude of low-frequency fluctuations in the regional BOLD signals) and the analogous indicators in the simulated signals, for each of the participants. With the obtained global influences, we reconstruct hidden excitability maps based on the pathology-dependent neuronal firing thresholds.

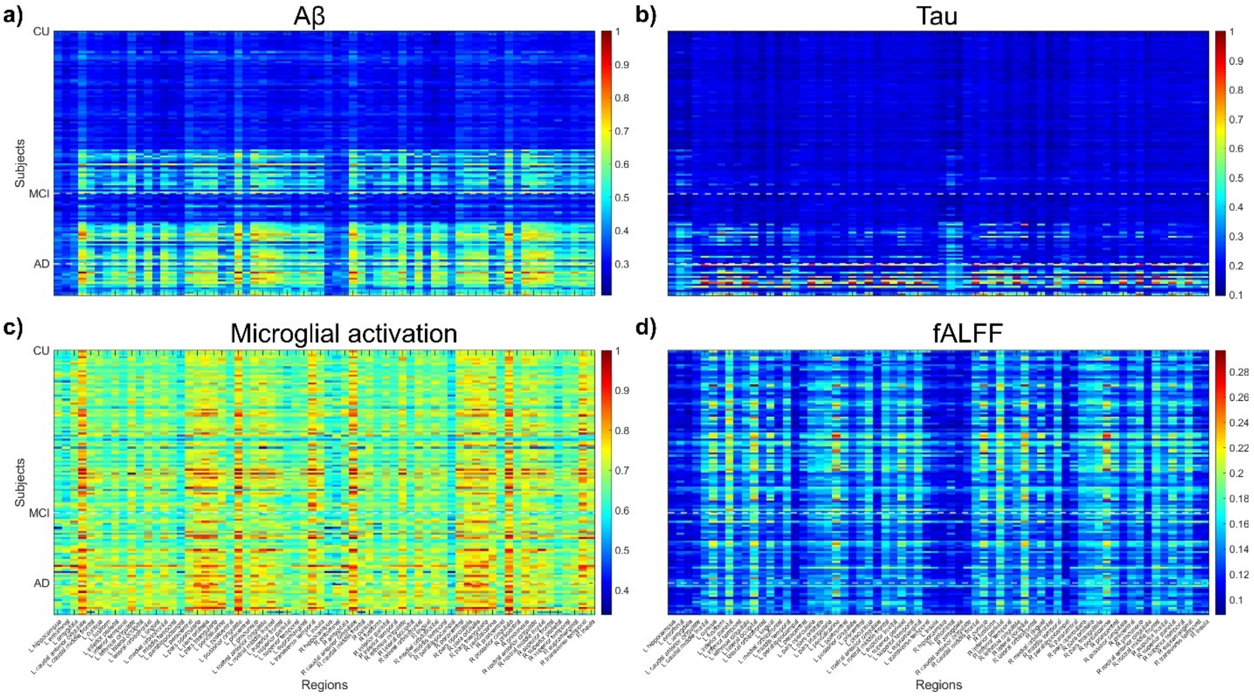


**Supplementary file 1—figure 7. Distributions of the processed data that was utilized in the study. a**) Aβ PET SUVRs **b**) Tau PET SUVRs **c**) Microglial activation PET SUVRs **d**) Fractional amplitudes of low-frequency fluctuations (fALFF) from fMRI. Subjects are organized by clinical groups (CU, MCI, AD) and appear in the same order as in Fig. 3 within a given clinical group. The SUVRs were normalized to the [0,1] interval by dividing by the absolute maximum value corresponding to each imaging modality.

**Supplementary file 1—table 1.** Demographics of the samples

| | CU | MCI | p | AD | p |
|---|---|---|---|---|---|
| Number of individuals (N) | 81 | 35 | - | 16 | - |
| Age (yrs), mean (s.d.) | 71.15 (7.73) | 72.17 (7.84) | 0.517 | 69.45 (9.10) | 0.438 |
| Female, N (%) | 63 (77.8) | 17 (48.6) | 0.004 | 8 (50.0) | 0.031 |
| Education (yrs), mean (s.d.) | 15.26 (3.54) | 15.63 (3.01) | 0.591 | 14.31 (3.38) | 0.327 |
| *APOE* ε4 carriers, N (%) | 24 (29.6) | 17 (48.6) | 0.059 | 7 (43.8) | 0.379 |
| MMSE, mean (s.d.) | 29.28 (0.97) | 28.40 (1.54) | < 0.001 | 20.25 (7.20) | < 0.001 |
| Aβ+, N (%) | 20 (24.7) | 23 (65.7) | 0.006 | 16 (100.0) | < 0.001 |

The reported p-values are for comparisons to cognitively unimpaired (CU) subjects. P-values for age, education and MMSE indicate values assessed with two-sided independent-samples t-tests. For the resting variables (sex, *APOE* ε4 status and Aβ+), Fischer exact test were performed. CU cognitively unimpaired; MCI mild cognitive impairment; AD Alzheimer's disease; *APOE* ε4, apolipoprotein epsilon 4; MMSE, Mini-Mental State examination.

**Supplementary file 1—table 2.** Summary demographics of the groups that were compiled for post-hoc analyses of the individually estimated pathophysiological quantities

| | CU(Aβ-) | MCI(Aβ-) | CU(Aβ+) | MCI(Aβ+) | AD(Aβ+) |
|---|---|---|---|---|---|
| Number of individuals (N) | 59 | 14 | 22 | 21 | 16 |
| Age (yrs), mean (s.d.) | 70.04 (8.29) | 72.89 (6.54) | 74.11 (5.03) | 71.68 (8.73) | 69.45 (9.10) |
| Female, N (%) | 45 (76.3) | 6 (42.9) | 18 (81.8) | 11 (52.4) | 8 (50.0) |
| Education (yrs), mean (s.d.) | 15.59 (3.79) | 14.86 (3.53) | 14.36 (2.61) | 16.14 (2.57) | 14.31 (3.38) |
| *APOE* ε4 carriers, N (%) | 13 (22.0) | 2 (14.3) | 11 (50.0) | 15 (71.4) | 7 (43.8) |
| MMSE, mean (s.d.) | 29.28 (0.96) | 28.64 (1.39) | 29.27 (1.03) | 28.24 (1.64) | 20.25 (7.20) |
| | Braak 0 | Braak I-II | Braak III-IV | Braak V-VI | |
| Number of individuals (N) | 66 | 33 | 18 | 15 | |
| Age (yrs), mean (s.d.) | 70.28 (8.01) | 72.89 (8.24) | 74.68 (5.09) | 67.44 (7.72) | |
| Female, N (%) | 45 (68.2) | 24 (72.7) | 10 (55.6) | 9 (60.0) | |
| Education (yrs), mean (s.d.) | 15.27 (3.61) | 15.03 (3.41) | 15.72 (2.42) | 15.00 (3.51) | |
| *APOE* ε4 carriers, N (%) | 16 (24.2) | 11 (33.3) | 14 (77.8) | 7 (46.7) | |
| MMSE, mean (s.d.) | 29.31 (0.89) | 28.24 (2.09) | 27.83 (2.81) | 21.47 (8.24) | |

CU cognitively unimpaired; MCI mild cognitive impairment; AD Alzheimer's disease; *APOE* ε4, apolipoprotein epsilon 4; MMSE, Mini-Mental State examination.

**Supplementary file 1—table 3.** Results of statistical tests

| Quantity | Test | Group 1 | Group 2 | F-Statistic | dfe | p |
|---|---|---|---|---|---|---|
| Ratio of power in theta band^ | ANCOVA post-hoc t-Tests | CU(Aβ-) | CU(Aβ+) | 41.88 | 77 | < 0.001 |
| | | CU(Aβ-) | MCI(Aβ+) | 54.63 | 76 | < 0.001 |
| | | CU(Aβ-) | AD(Aβ+) | 41.84 | 71 | < 0.001 |
| | | MCI(Aβ-) | CU(Aβ+) | 6.50 | 32 | 0.016 |
| | | MCI(Aβ-) | MCI(Aβ+) | 21.28 | 31 | < 0.001 |
| | | MCI(Aβ-) | AD(Aβ+) | 17.70 | 26 | < 0.001 |
| | | Braak 0 | Braak III-IV | 14.34 | 80 | < 0.001 |
| | | Braak 0 | Braak V-VI | 33.60 | 77 | < 0.001 |
| | | Braak I-II | Braak V-VI | 8.23 | 44 | 0.006 |
| Ratio of power in alpha1 band^ | ANCOVA post-hoc t-Tests | CU(Aβ-) | CU(Aβ+) | 31.78 | 77 | < 0.001 |
| | | CU(Aβ-) | MCI(Aβ+) | 33.40 | 76 | < 0.001 |
| | | CU(Aβ-) | AD(Aβ+) | 20.12 | 71 | < 0.001 |
| | | MCI(Aβ-) | MCI(Aβ+) | 12.14 | 31 | 0.001 |
| | | MCI(Aβ-) | AD(Aβ+) | 7.36 | 26 | 0.011 |
| | | Braak 0 | Braak III-IV | 5.96 | 80 | 0.017 |
| | | Braak 0 | Braak V-VI | 16.95 | 77 | < 0.001 |
| | | Braak I-II | Braak V-VI | 8.26 | 44 | 0.006 |
| Mean excitatory activity^ | ANCOVA post-hoc t-Tests | CU(Aβ-) | CU(Aβ+) | 49.80 | 77 | < 0.001 |
| | | CU(Aβ-) | MCI(Aβ+) | 59.34 | 76 | < 0.001 |
| | | CU(Aβ-) | AD(Aβ+) | 46.93 | 71 | < 0.001 |
| | | MCI(Aβ-) | CU(Aβ+) | 8.86 | 32 | 0.005 |
| | | MCI(Aβ-) | MCI(Aβ+) | 21.25 | 31 | < 0.001 |
| | | MCI(Aβ-) | AD(Aβ+) | 18.22 | 26 | < 0.001 |
| | | Braak 0 | Braak III-IV | 15.35 | 80 | < 0.001 |
| | | Braak 0 | Braak V-VI | 38.38 | 77 | < 0.001 |
| | | Braak I-II | Braak V-VI | 8.70 | 44 | 0.005 |
| Average intra-brain excitability^ | ANCOVA post-hoc t-Tests | CU(Aβ-) | CU(Aβ+) | 55.54 | 77 | < 0.001 |
| | | CU(Aβ-) | MCI(Aβ+) | 66.06 | 76 | < 0.001 |
| | | CU(Aβ-) | AD(Aβ+) | 54.19 | 71 | < 0.001 |
| | | MCI(Aβ-) | CU(Aβ+) | 9.96 | 32 | 0.003 |
| | | MCI(Aβ-) | MCI(Aβ+) | 22.65 | 31 | < 0.001 |
| | | MCI(Aβ-) | AD(Aβ+) | 19.78 | 26 | < 0.001 |
| | | Braak 0 | Braak I-II | 4.48 | 95 | 0.037 |

| | | | | | | |
|---|---|---|---|---|---|---|
| | | Braak 0 | Braak III-IV | 18.02 | 80 | < 0.001 |
| | | Braak 0 | Braak V-VI | 44.35 | 77 | < 0.001 |
| | | Braak I-II | Braak V-VI | 8.97 | 44 | 0.004 |
| Plasma biomarkers | Spearman correlation | Mean excitability | p-tau181 (*r=0.39*) | NA | NA | 0.003 |
| | | Mean excitability | p-tau231 (*r=0.38*) | NA | NA | 0.003 |
| | | Mean excitability | p-tau217 (*r=0.52*) | NA | NA | < 0.001 |
| | | Mean excitability | GFAP (*r=0.32*) | NA | NA | < 0.001 |
| MMSE* | Linear regression model | NA | NA | 4.43 | 123 | < 0.001 |
| MoCA* | Linear regression model | NA | NA | 4.58 | 120 | < 0.001 |

CU cognitively unimpaired; MCI mild cognitive impairment; AD Alzheimer's disease; ^ sex and age adjusted; * sex, age and education adjusted.

**Supplementary file 1—table 4.** Multiple linear regression analysis investigating the pathological effects on neuronal activity as predictors of MMSE and MoCA scores in the *Aβ, tau, Aβ·tau, microglial activation* influence model

| | **MMSE scores** | | |
|---|---|---|---|
| | β | 95% CI of β | p |
| Intercept | 27.136 | [21.110 34.161] | < 0.001 |
| $\theta^{A\beta}$ | 0.373 | [-0.296 1.041] | 0.272 |
| $\theta^{Tau}$ | 0.151 | [-0.524 0.837] | 0.659 |
| **$\theta^{A\beta\cdot Tau}$** | **1.530** | **[0.856 2.205]** | **< 0.001** |
| $\theta^{\text{microglial act.}}$ | 45 | [-1.1402 0.249] | 0.201 |
| Sex | -0.960 | [-2.348 0.429] | 0.174 |
| Age | 0.009 | [-0.080 0.099] | 0.834 |
| Education | 0.029 | [-0.163 0.221] | 0.762 |
| | **MoCA scores** | | |
| | β | 95% CI of β | p |
| Intercept | 22.145 | [11.994 32.296] | < 0.001 |
| **$\theta^{A\beta}$** | **1.004** | **[0.081 1.927]** | **0.033** |
| $\theta^{Tau}$ | 0.129 | [-0.791 1.048] | 0.782 |
| **$\theta^{A\beta\cdot Tau}$** | **1.966** | **[1.024 2.888]** | **< 0.001** |
| $\theta^{\text{microglial act.}}$ | -0.609 | [-1.565 0.347] | 0.209 |
| Sex | -1.624 | [-3.570 0.322] | 0.101 |
| Age | 0.058 | [-0.073 0.189] | 0.380 |
| Education | -0.025 | [-0.2920 0.2424] | 0.855 |

The influences of Aβ plaques ($\theta_E^{A\beta}$), tau tangles ($\theta_E^{Tau}$), the interaction of Aβ and tau ($\theta_E^{A\beta\cdot Tau}$) and microglial activation ($\theta_E^{microglial\ act.}$) on neuronal activity, sex, age and education were considered as predictors. Reported values are obtained coefficients ($\beta$), the 95% confidence intervals and the p-values for the t-statistic of the two-sided hypothesis tests. Significant terms (5% level) other than the intercepts are highlighted. MMSE: $R^2$=0.19, $p < 0.001$; MoCA: $R^2$=0.21, $p < 0.001$. MMSE, Mini-Mental State examination; MoCA, Montreal Cognitive Assessment.

**Supplementary file 1—table 5.** Brain regions in the considered parcellation.

| | | |
|---|---|---|
| Hippocampus | Lateral Orbitofrontal[B] | Posterior Cingulate |
| Entorhinal[R] | Medial Orbitofrontal[R] | Precentral |
| Amygdala | Lingual[L] | Precuneus[B] |
| Caudal Anterior Cingulate | Middle Temporal[B] | Rostral Anterior Cingulate |
| Caudal Middle Frontal | Parahippocampal[B] | Rostral Middle Frontal |
| Cuneus | Paracentral | Superior Frontal |
| Fusiform[B] | Pars Opercularis[R] | Superior Parietal |
| Inferior Parietal[R] | Pars Orbitalis[2] | Superior Temporal |
| Inferior Temporal[B] | Pars Triangularis[R] | Supramarginal |
| Isthmus Cingulate[L] | Pericalcarine[L] | Transverse Temporal |
| Lateral Occipital[B] | Postcentral | Insula |

Regions with a superscript next to the name have statistically different microglial activation PET SUVRs when comparing the CU and AD groups: [B] - bilaterally different, [L] - only the values of the left hemisphere region is different, [R] - only the values of the right hemisphere region is different.

**Supplementary file 1—table 6.** Electrophysiological model parameters

| Parameter | Definition | Value | Ref. |
|---|---|---|---|
| $\tau_I$ | Time-constant controlling the decay of inhibitory activity after stimulation | 0.02 s | (Abeysuriya et al., 2018) |
| $\tau_E$ | Time-constant controlling the decay of excitatory activity after stimulation | 0.01 s | (Abeysuriya et al., 2018) |
| $C_{II}$ | Local inhibitory-inhibitory connection strength | 1.2 | (Gjorgjieva et al., 2016; Meijer et al., 2015; Wilson & Cowan, 1972) |
| $C_{EI}$ | Local excitatory-inhibitory connection strength | 6 | (Gjorgjieva et al., 2016; Meijer et al., 2015; Wilson & Cowan, 1972) |
| $C_{EE}$ | Local excitatory-excitatory connection strength | 6.4 | (Gjorgjieva et al., 2016; Meijer et al., 2015; Wilson & Cowan, 1972) |
| $C_{IE}$ | Local inhibitory-excitatory connection strength | 4.8 | (Gjorgjieva et al., 2016; Meijer et al., 2015; Wilson & Cowan, 1972) |
| $P$ | Average constant external input received by the excitatory population | 0.65<br>(set to produce plausible simulated electrophysiological and BOLD signals) | (Gjorgjieva et al., 2016; Meijer et al., 2015; Wilson & Cowan, 1972) |
| $a_I$ | Maximum slope of the inhibitory sigmoidal activation function | 1 | (Abeysuriya et al., 2018) |
| $a_E$ | Maximum slope of the excitatory sigmoidal activation function | 1 | (Abeysuriya et al., 2018) |
| $\theta_I$ | Position of the inhibitory sigmoidal firing function' threshold for activation | 4 | (Gjorgjieva et al., 2016; Meijer et al., 2015; Wilson & Cowan, 1972) |
| $\theta_E$ | Position of the excitatory sigmoidal firing function' threshold for activation | Variable in [2.75,2.85] depending on the regional pathological loads<br><br>2.8 (in normal baseline conditions) | (Abeysuriya et al., 2018; Daffertshofer & van Wijk, 2011; Gjorgjieva et al., 2016; Meijer et al., 2015; Wilson & Cowan, 1972) |
| $\eta$ | Global coupling strength scaling the anatomical connectivity matrix $C_{lk}$ | 2<br>(set to produce plausible simulated electrophysiological and BOLD signals) | (Abeysuriya et al., 2018; Daffertshofer & van Wijk, 2011; Gjorgjieva et al., 2016; Meijer et |

| | | | |
|---|---|---|---|
| | | | al., 2015; Wilson & Cowan, 1972) |
| $N$ | Number of brain regions of interest | 66 | (Klein & Tourville, 2012) |

**Supplementary file 1—table 7.** Metabolic/hemodynamic model parameters

| Parameter | Definition | Value | Ref. |
|---|---|---|---|
| $h_E$ | Efficacy of glucose consumption response to excitation | 1 | (Sotero et al., 2009; Sotero & Trujillo-Barreto, 2007, 2008; Valdes-Sosa et al., 2009) |
| $h_I$ | Efficacy of glucose consumption response to inhibition | 1 | (Sotero et al., 2009; Sotero & Trujillo-Barreto, 2007, 2008; Valdes-Sosa et al., 2009) |
| $\varkappa_E$ | Time-constant of the excitatory glucose consumption impulse response. | 1 s | (Sotero et al., 2009; Sotero & Trujillo-Barreto, 2007, 2008; Valdes-Sosa et al., 2009) |
| $\varkappa_I$ | Time-constant of the inhibitory glucose consumption impulse response. | 1 s | (Sotero et al., 2009; Sotero & Trujillo-Barreto, 2007, 2008; Valdes-Sosa et al., 2009) |
| $c$ | Steepness of the sigmoid function $x$ | 2.5 | (Sotero et al., 2009; Sotero & Trujillo-Barreto, 2007, 2008; Valdes-Sosa et al., 2009) |
| $d$ | Position of the threshold of the sigmoid function $x$ | 1.6 | (Sotero et al., 2009; Sotero & Trujillo-Barreto, 2007, 2008; Valdes-Sosa et al., 2009) |
| $\gamma$ | Baseline ratio of excitatory to inhibitory synaptic activity in the voxel | 5 | (Sotero et al., 2009; Sotero & Trujillo-Barreto, 2007, 2008; Valdes-Sosa et al., 2009) |
| $x_0$ | Fraction of glucose following the glycogenolitic pathway at rest | $\frac{1}{1+exp\left[c\left(d-1(t)\right)\right]}$ | (Sotero et al., 2009; Sotero & Trujillo-Barreto, 2007, 2008; Valdes-Sosa et al., 2009) |
| $\mu$ | Efficacy of blood flow response to excitation | 0.8 | (Sotero et al., 2009; Sotero & Trujillo-Barreto, 2007, 2008; Valdes-Sosa et al., 2009) |
| $\varkappa_f$ | Time constant for CBF response | 1.7 | (Sotero et al., 2009; Sotero & Trujillo-Barreto, 2007, |

| | | | |
|---|---|---|---|
| | | | 2008; Valdes-Sosa et al., 2009) |
| $\varkappa_0$ | Transit time through the balloon | 1 | (Sotero et al., 2009; Sotero & Trujillo-Barreto, 2007, 2008; Valdes-Sosa et al., 2009) |
| $\zeta$ | Coefficient of the steady state flow-volume relationship | 0.4 | (Sotero et al., 2009; Sotero & Trujillo-Barreto, 2007, 2008; Valdes-Sosa et al., 2009) |
| $V_0$ | Baseline blood volume | 0.03 | (Sotero et al., 2009; Sotero & Trujillo-Barreto, 2007, 2008; Valdes-Sosa et al., 2009) |
| $\Upsilon_0$ | frequency offset of a fully deoxygenated blood vessel at 3 T | 80.6 $s^{-1}$ (at 3 T) | (Archila-Meléndez et al., 2020; Obata et al., 2004; Simon & Buxton, 2015) |
| $r_0$ | Slope defining the dependence of the R2*relaxation rate on blood oxygenation | 178 $s^{-1}$ (at 3 T) | (Archila-Meléndez et al., 2020; Obata et al., 2004; Simon & Buxton, 2015) |
| $E_0$ | Baseline oxygen extraction fraction | 0.4 | (Archila-Meléndez et al., 2020; Obata et al., 2004; Simon & Buxton, 2015) |
| $\varepsilon$ | Intrinsic ratio of blood to tissue signals at rest | 0.24 | (Archila-Meléndez et al., 2020; Obata et al., 2004; Simon & Buxton, 2015) |
| $TE$ | Echo time | 32.0 ms | https://triad.tnl-mcgill.com/ |

**Supplementary file 1—pseudocode 1.**

```
// program to calculate the personalized combined neuronal activity influences by Aβ, tau and
// Aβ · Tau

{       // definitions
Define surrogate optimization parameters
Load the subject's Aβ, tau and fALFF (rs-fMRI) and anatomical connectivity matrix
Define the neuronal activity influence model (Eq. 1)
Define neural mass model and transformations to simulate the resting-state BOLD signal
Define the objective function (minimizes distance between real and simulated BOLD)
}

{       // optimization
FOR i = 1 TO 20         // different random optimization evaluation trials
        Perform surrogate optimization until the algorithm converges
                // At each iteration:
                // simulate the BOLD signal,
                // calculate similarity with the subject's real signal,
                // retain the best evaluation thus far
                // (performed by Matlab's surrogateopt.m)
        Save the optimized neuronal activity affectation parameters and optimization outputs
ENDFOR
}

{       // post-processing
Retain the optimization outcome with the lowest overall cost
Reconstruct hidden quantities of interest, e.g., neuronal excitabilities, spectral power, etc
}
```

### Supplementary file 1—References


Abeysuriya, R. G., Hadida, J., Sotiropoulos, S. N., Jbabdi, S., Becker, R., Hunt, B. A. E., Brookes, M. J., & Woolrich, W. (2018). *A biophysical model of dynamic balancing of excitation and inhibition in fast oscillatory large-scale networks*.
